# Brain-behavior patterns define a dimensional biotype in medication-naïve adults with attention-deficit hyperactivity disorder

**DOI:** 10.1101/190660

**Authors:** Hsiang-Yuan Lin, Luca Cocchi, Andrew Zalesky, Jinglei Lv, Alistair Perry, Wen-Yih Isaac Tseng, Prantik Kundu, Michael Breakspear, Susan Shur-Fen Gau

## Abstract

**Background:** Childhood-onset attention-deficit hyperactivity disorder (ADHD) in adults is clinically heterogeneous and commonly presents with different patterns of cognitive deficits. It is unclear if this clinical heterogeneity expresses a dimensional or categorical difference in ADHD.

**Methods:** We first studied differences in functional connectivity in multi-echo resting-state functional magnetic resonance imaging (rs-fMRI) acquired from 80 medication-naïve adults with ADHD and 123 matched healthy controls. We then used canonical correlation analysis (CCA) to identify latent relationships between symptoms and patterns of altered functional connectivity (dimensional biotype) in patients. Clustering methods were implemented to test if the individual associations between resting-state brain connectivity and symptoms reflected a non-overlapping categorical biotype.

**Results:** Adults with ADHD showed stronger functional connectivity compared to healthy controls, predominantly between the default-mode, cingulo-opercular and subcortical networks. CCA identified a single mode of brain-symptom co-variation, corresponding to an ADHD dimensional biotype. This dimensional biotype is characterized by a unique combination of altered connectivity correlating with symptoms of hyperactivity-impulsivity, inattention, and intelligence. Clustering analyses did not support the existence of distinct categorical biotypes of adult ADHD.

**Conclusions:** Overall, our data advance a novel finding that the reduced functional segregation between default-mode and cognitive control networks supports a clinically important dimensional biotype of childhood-onset adult ADHD. Despite the heterogeneity of its presentation, our work suggests that childhood-onset adult ADHD is a single disorder characterized by dimensional brain-symptom mediators.

## Introduction

Attention-deficit hyperactivity disorder (ADHD) is a childhood-onset neurodevelopmental disorder whose persistence into adulthood has been increasingly acknowledged (Asherson *et al.*, 2016). ADHD in both children and adults manifests as a heterogeneous condition with significantly varied intensity and types of inattention and hyperactivity-impulsivity symptoms across individuals (Asherson *et al.*, 2016). Individuals with ADHD also commonly present with general cognitive deficits (Fair *et al.*, 2012; Mostert *et al.*, 2015) interacting with clinical symptoms (Cheung *et al.*, 2015; Rommelse *et al.*, 2016).

Although the aetiological markers of ADHD remain elusive, the disorder is associated with functional alterations in whole-brain resting-state networks (Gallo and Posner, 2016). ADHD symptoms, in both adults and children, have consistently been linked to abnormal functional connectivity within and between the default-mode network (DMN), cognitive control and attention networks, as well as mesocorticolimbic circuits (Cocchi *et al.*, 2012; Lin and Gau, 2015; Castellanos and Aoki, 2016; Cai *et al.*, 2017). In line with the clinical expression of ADHD, these networks are widely-known to provide crucial support to higher-order cognitive functions (Cole *et al.*, 2016) and general intelligence (Hearne *et al.*, 2016).

Altered resting-state connectivity has been used to parse clinically heterogeneous children with ADHD into more coherent subgroups (Gates *et al.*, 2014; Costa Dias *et al.*, 2015). However, attempts to stratify ADHD based on resting-state connectivity have so far provided inconsistent results with important caveats from a clinical perspective. For example, connectivity-based clusters have not converged on symptomatically similar subgroups of ADHD across studies (Gates *et al.*, 2014). Previous studies have also isolated subgroups by pooling ADHD and typically developing controls together, precluding a distinct assessment of possible categories specific to ADHD (Gates *et al.*, 2014; Costa Dias *et al.*, 2015; Marquand *et al.*, 2016). On the other hand, dimensional models of ADHD psychopathology based on behavioral measures are increasingly supported (Marcus and Barry, 2011; Willcutt *et al.*, 2012). Nevertheless, the link between existing dimensional models and brain function remains elusive, with very limited data in the adult population (Asherson *et al.*, 2016; Gallo and Posner, 2016).

Recent advances in neuroimaging methods provide new opportunities to test the dimensional and categorical models of ADHD by using multivariate analyses (e.g., canonical correlation analysis, CCA) of brain-behavior relationships (Smith *et al.*, 2015; Drysdale *et al.*, 2017). Leveraging these new methodological developments, the heterogeneity between individuals with ADHD can be addressed by borrowing and extending the construct of biotype. Here, a biotype delineates a subgroup of individuals with ADHD sharing common symptom patterns and abnormal resting-state connectivity. A dimensional biotype is defined by abnormal brain-symptom patterns that vary in severity across individuals on a continuum. Conversely, a categorical biotype defines a discrete subgroup of individuals sharing a combined pattern of altered functional connectivity and clinical-cognitive features that are unique and segregated from other individuals. Breaking down ADHD heterogeneity into dimensional or categorical biotypes is important because it will enable the development of targeted research and clinical interventions. Prior studies have attempted to define ADHD categorical subtypes based on co-occurring psychiatric symptoms (Acosta *et al.*, 2008), neuropsychological profiles (Fair *et al.*, 2012; Mostert *et al.*, 2015), temperaments (Karalunas *et al.* 2014), and patterns of functional connectivity (Gates *et al.*, 2014; Costa Dias *et al.*, 2015). Karalunas *et al.* (2014) were the first to use behavioral and physiological measures to investigate the existence of ADHD categorical subtypes. In this study, subtypes were initially defined using temperamental features and subsequently validated by cardiac and brain measures, as well as clinical outcomes. However, to the best of our knowledge, no studies have combined symptoms and brain features to assess the existence of categorical and/or dimensional biotypes of ADHD. In other psychiatric disorders such as psychosis, neuroimaging and behavioral data have recently been used to parse individuals into separate biotypes. For example, Clementz *et al.* (2016) delineated three categorical psychosis biotypes using cluster analysis applied to electrophysiological and neuropsychological data, while Kaczkurkin *et al.* (2017) applied a constrained factor analysis to psychiatric symptoms in a large cohort of youth to delineate orthogonal dimensions of psychopathology.

In this study, we assessed complex relationships between clinical and cognitive presentations and abnormal resting-state connectivity in a large medication-naïve sample of adults with ADHD. The principal aim of our analysis was to uncover the existence of dimensional or/and categorical biotypes of childhood-onset adult ADHD. Specifically, we identified distinct alterations in whole-brain resting-state connectivity (as estimated by a multi-echo planar imaging, EPI, acquisition) that were associated with distinct symptoms of inattention and hyperactivity-impulsivity, as well as general cognitive functioning (Wechsler, 1997). To maximize the translational value of our work (Drysdale *et al.*, 2017), we focused on a subset of symptom measures which capture the majority of variance in ADHD phenomenology (Asherson *et al.*, 2016) and are commonly acquired in clinical settings. Based on recent findings (Barch, 2017; Drysdale *et al.*, 2017), we hypothesized that biotypes of adult ADHD would involve functional connectivity changes in brain networks previously with symptoms of ADHD. Specifically, we expected the interaction between the DMN and cognitive control and attention networks to be key to delineation of ADHD dimensional or categorical biotypes (Cocchi *et al.*, 2012; Lin and Gau, 2015; Cai *et al.*, 2017).

## Methods

### Participants and procedure

The study included 80 medication-naïve adults with childhood-onset ADHD (24 females) aged 18-39 years (mean 26.7 years), who fulfilled *DSM-IV-TR* criteria for ADHD, and 123 age-(mean 25.7 years) sex-(44 females) and IQ-matched healthy controls with no clinically significant psychopathology. The participants were assessed at the adult ADHD special clinic of the Department of Psychiatry, National Taiwan University Hospital (NTUH), Taipei, Taiwan from March 2014 to December 2016. Compared to existing multi-sites studies (Hoogman *et al.*, 2017), we recruited a homogeneous sample that is free from medication and multi-site confounds (e.g., different MR scanners and protocols). The detection of biotypes in our unique sample is also facilitated because the variance in the data cannot be linked to heterogeneous developmental delays and psychiatric comorbidity (Schnack and Kahn, 2016). From a methodological viewpoint, our analyses were tailored so that meaningful ADHD biotypes could be detected with the given sample size (see Supplementary Methods).

All participants, including patients and healthy controls, were recruited via advertisements at the hospital, colleges, and websites. Adults interested in the study were first telephone-screened using the Chinese version of the Adult ADHD Self-Report Scale v1.1 (Yeh *et al.*, 2008). Those identified as probable cases of ADHD by telephone interview were excluded from the study if they had ever received psychotropic treatment before. All other participants were then clinically assessed and diagnosed by a board-certified child psychiatrist (SS Gau) for the presence or absence of ADHD and any other psychiatric disorders. The ADHD diagnosis was further confirmed with the Conners’ Adult ADHD Diagnostic Interview (Conners *et al.*, 1999) for current ADHD, and the modified adult version of the ADHD supplement of the Chinese version of the Kiddie-Schedule for Affective Disorders and Schizophrenia-Epidemiological Version (K-SADS-E) for childhood and current ADHD (see Supplementary Information) (Lin *et al.*, 2016). Age-matched healthy adult controls without any lifetime diagnosis of ADHD received the same clinical assessments as the ADHD group. Exclusion criteria for all participants included: Any prior systemic medical illness; a history of affective disorders, psychosis, substance use disorder, autism spectrum disorder; current depressive/anxiety symptoms or suicidal ideation; a history of psychotropic treatment, including medication for ADHD; and a full IQ (FIQ) <80, as assessed by Wechsler Adult Intelligence Scale-3^rd^ Edition (Wechsler, 1997). Of the ADHD participants, 47, 32, and one were diagnosed with the DSM-IV inattentive, combined, and hyperactivity-impulsive subtypes, respectively (Table 1 and Supplementary Table 1). Notably, only 61 adults retained the same subtypes across current and childhood presentations (the childhood subtypes were ascertained using the K-SADS-E). All study procedures were approved by the Research Ethics Committee of the NTUH (201401024RINC; ClinicalTrials.gov number, NCT02642068). All participants provided written informed consent.

**Table 1.** Demographic and clinical features of the participants

| Mean (SD) | Control (N=123) | ADHD (N=80) | Statistics |
| --- | --- | --- | --- |
| Age (18-39 years) | 25.7 (4.9) | 26.7 (5.6) | $p=0.171$ |
| Sex (M/F) | 79/44 | 56/24 | $p=0.394$ |
| Handedness (R/L) | 118/5 | 63/16 | $p<0.001$ |
| FIQ | 109.8 (9.3)<br>(range: 89-138) | 107.5 (10.3)<br>(range: 80-137) | $p=0.096$ |
| VIQ | 108.2 (9.1) | 105.9 (11.2) | $p=0.120$ |
| PIQ | 110.6 (11.4) | 108.4 (16.1) | $p=0.268$ |
| *ADHD symptoms* |  |  |  |
| Inattention<sup>a</sup> | 6.5 (4.8) | 19.7 (5.0) | $p<0.001$ |
| Hyperactivity/Impulsivity<sup>a</sup> | 3.1 (4.3) | 13.6 (6.5) | $p<0.001$ |
| Opposition-defiance<sup>a</sup> | 3.6 (11.4) | 4.2 (5.9) | $p<0.001$ |
| ASRS-A | 13.2 (5.1) | 27.1 (4.8) | $p<0.001$ |
| ASRS-B | 8.9 (5.0) | 19.9 (6.22) | $p<0.001$ |
| Mean frame-wise displacement<sup>b</sup><br>(mm) | 0.045 (0.020)<br>(range: 0.014-0.123) | 0.049 (0.025)<br>(range: 0.017-0.108) | $p=0.233$ |

<sup>a</sup> Measured by the Swanson, Nolan, and Pelham, version IV (SNAP-IV) scale.
<sup>b</sup> Estimated by the Euclidian norm (enorm: square root of the sum of squares of the differences in motion derivatives), computed with AFNI's 1d\_tool.py.
Abbreviation: ADHD=attention-deficit hyperactivity disorder; FIQ=full intelligence quotient; PIQ=performance intelligence quotient; VIQ=verbal intelligence quotient; ASRS=Adult ADHD Self-Report Scale; M=male; F=female; R=right; L=left; SD=standard deviation.

### Imaging protocols and preprocessing

Neuroimaging data were acquired with a Siemens 3T Tim Trio scanner using a 32-channel head coil at Advanced Biomedical Imaging Lab of the NTUH. All participants were instructed to relax, clear their mind, and remain still with their eyes closed while undergoing a 7 min 39 sec resting-state functional magnetic resonance imaging (rs-fMRI) scan. Wakefulness was checked immediately after the sequence was completed, and all participants denied falling asleep during rs-fMRI scans. The duration of the rs-fMRI sequence was decided to limit the chance of possible fluctuations in participants’ wakefulness and in-scanner motion while allowing the acquisition of sufficient blood-oxygen-level-dependent (BOLD) signal (Van Dijk *et al.*, 2010; Birn *et al.*, 2013). Localizer and rs-fMRI scans were obtained first. T1-weighted anatomical scans (MPRAGE pulse sequences) and diffusion spectrum imaging, which was not analyzed here, were acquired thereafter. Functional images were acquired with a multi-echo EPI sequence (repetition time=2.55 sec; flip angle=90°; matrix size=64 × 64; in-plane resolution=3.75 mm; F0V=240 mm; 31 oblique slices, alternating slice acquisition slice thickness 3.75 mm with 10% gap; iPAT factor=3; bandwidth=1,698 Hz/pixel; echo time, TE=12, 28, 44 and 60 msec). Anatomical images were acquired using a T1-weighted magnetization prepared rapid gradient echo (MPRAGE) sequence (repetition time=2 sec; TE=2.98 msec; flip angle=9°; matrix size=256×256; inversion time=900 msec; voxel size=1 mm^3^).

These fMRI data permit unique denoising and analyses using multi-echo independent components analysis (ME-ICA v3.0 beta1; www.bitbucket.org/prantikk/me-ica) (Kundu *et al.*, 2012). In comparison to standard single-echo imaging, this approach enhances the signal-to-noise ratio and addresses concerns regarding non-neural confounding factors in fMRI acquisitions, particularly head motion (Kundu *et al.*, 2017). Preprocessing was carried out using AFNI v16.1.10 and Python v2.7.11 toolkits. Each individual’s denoised EPI was co-registered to their T1 image and then non-linearly normalized to the Montreal Neurological Institute template (3-mm isotropic voxel size). Spatial smoothing was not conducted for the denoised normalized data as per the recommendation by Lombardo *et al.* (2016). The time course was temporally band-pass filtered (0.01~0.1 Hz). *Post-hoc* analysis showed that levels of framewise displacement (computed using AFNI’s 1d_tool.py) did not significantly differ between ADHD and controls (*F*(1, 201)=1.462, p=0.233; Table 1). No participants exhibited extreme levels of head motion (translation and rotation rigid-body realignment estimates were <1.5mm and <1.5°). ME-ICA are detailed in Supplementary Methods.

### Whole-brain patterns of altered functional connectivity in ADHD

We delineated ADHD biotypes based on distinct patterns of abnormal resting-state connectivity identified by diagnostic group comparisons (ADHD vs. controls). We reasoned that biologically meaningful ADHD subtypes would be best characterized by a subset of the whole-brain resting-state connectivity features that comprise group differences between ADHD and controls. A graphic representation of the analysis pipeline is presented in Fig. 1.

**Figure 1.**
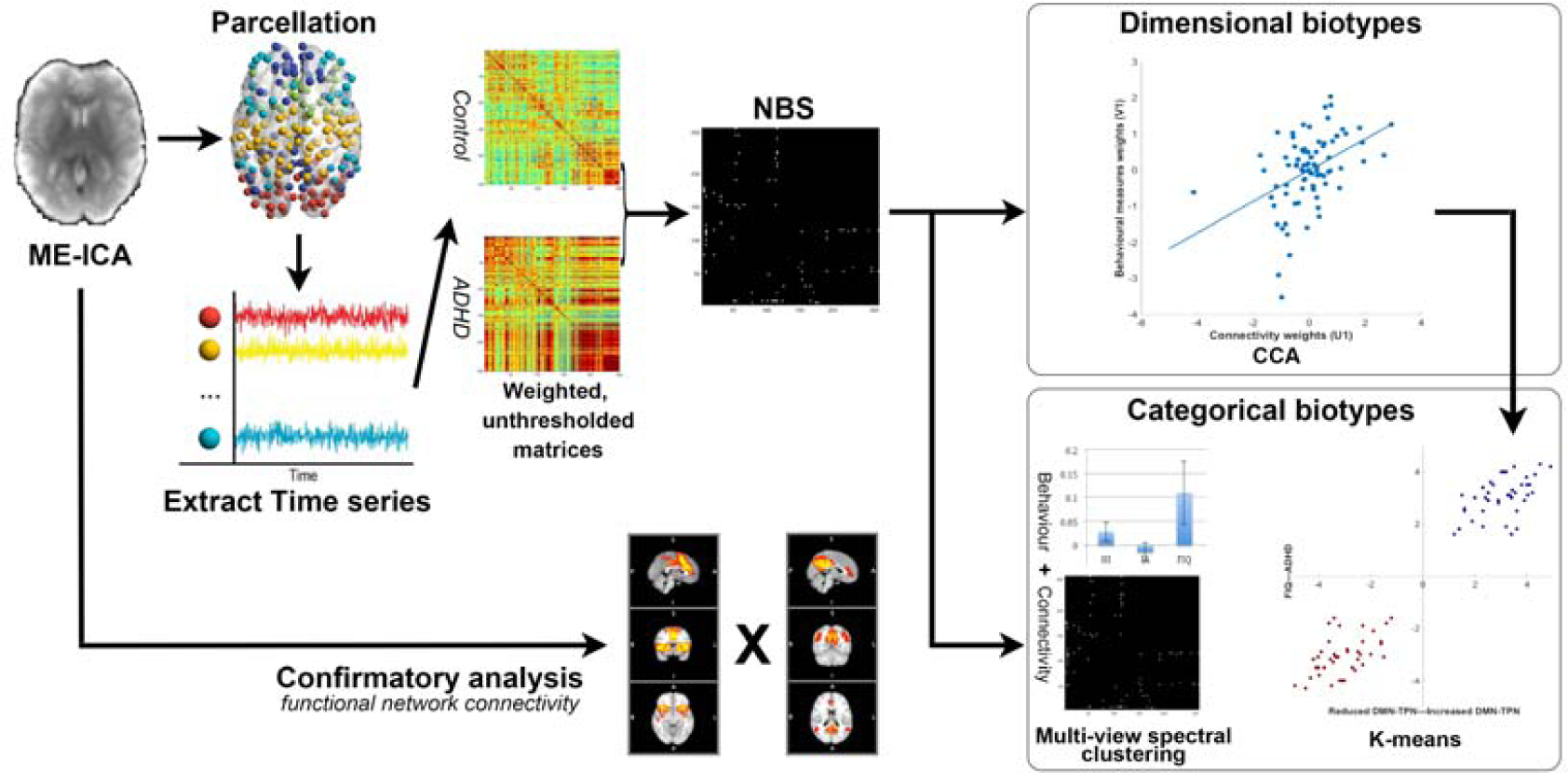
Overview of data processing and analysis pipeline. Multi-echo independent component analysis (ME-ICA) was used to denoise eye-closed resting-state imaging data. Functional connectivity matrices were calculated by extracting the average signal time series from validated brain parcellations. The network-based statistic (NBS) was used to assess differences in whole-brain connectivity between 80 drug-naive individuals with ADHD and 123 matched healthy controls. Results from the NBS were corroborated by a data-driven functional network connectivity analysis. Following data reduction steps (see text) functional connectivity and behavioral data were used to perform a canonical correlation analysis (CCA). This analysis allowed to identify hidden relations between resting-state functional connectivity and behavior. The CCA assigns a weight for each functional connection and each symptom such that the level of linear correlation between connectivity and behavioral data across individuals with ADHD is maximized. Permutation testing determined a family-wise error (FWE) corrected p-value for this multivariate correlation (i.e., dimensional biotype). Regarding the discovery of categorical biotype, two complementary approaches were employed: a. *k*-means clustering was used to test if the individual associations between connectivity and behavior form distinct clusters; b. multi-view spectral clustering (Kumar and Daumé, 2011) was implemented using features from symptoms and connectivity (NBS results).

We generated a whole-brain correlation matrix for each participant, based on a well-validated functional brain parcellation (Power *et al.*, 2011). Eleven of the originally published 264 nodes (mostly the inferior temporal areas and cerebellum areas, including 2 regions-of-interest, ROIs, in the DMN; 5 ROIs in the limbic network; 4 ROIs in the cerebellum) were excluded from analysis owing to incomplete MRI coverage. We additionally included the bilateral nucleus accumbens, centered at MNI coordinates x=±9,y=9,z=-8 as defined by Costa Dias *et al.* (2015). These ROIs were included for their roles implicated in ADHD. The final parcellation included 255 functional nodes (10-mm-diameter spheres). fMRI time series were extracted from each node by averaging over the representative voxels. The Fisher z-transformation was applied to each correlation coefficient within the matrix.

We employed network-based statistic (NBS) (Zalesky *et al.*, 2010), a validated non-parametric algorithm that controls for multiple comparisons (between all possible edges). NBS was performed on connectivity matrices to identify bivariate patterns of connectivity that differentiated ADHD from healthy controls. NBS is based on the principles underpinning traditional cluster-based thresholding of statistical parametric maps and hence proceeds with a preliminary height threshold (pair-wise connections) followed by a familywise error (FWE)-corrected cluster threshold (topological subnetworks of connections). We used a t-statistic (absolute value) of 3.5 as our initial height threshold (corresponding to p<0.0005, two-tailed), followed by a conservative FWE corrected p<0.05 for our pair-wise contrasts. This height threshold was decided based on our previous neuroimaging investigation in adult ADHD (Cocchi *et al.*, 2012). The inference was performed using permutation testing (10,000 permutations).

To ensure our results are not specific to any particular parcellation template, we repeated the analysis with functional networks constructed using parcellation templates of different resolution and nature (Supplementary Fig. 1). To further test the robustness of our NBS results, we undertook a confirmatory analysis using data-driven group ICA and functional network connectivity (Jafri *et al.*, 2008) (see Fig. 1 and Supplementary Information).

### CCA and clustering

CCA is a multivariate statistical method to identify latent, linear relations between combinations of independent and dependent variables (Krzanowski, 2000). The first mode represents canonical correlations corresponding to the maximum co-variation between the two sets of variables. Subsequent modes represent maximum residual, orthogonal co-variation. We implemented CCA, in steps similar to those previously reported (Smith *et al.*, 2015; Perry *et al.*, 2017), to identify modes which relate sets of resting-state connectivity patterns to symptom and cognitive measures of ADHD pathology, within the ADHD cohort. Connections composing the altered resting-state network as identified by the NBS were selected as connectivity features within the CCA. Three symptom measures were included in the CCA: factor scores of inattentive and hyperactivity-impulsivity symptoms, and FIQ. As general cognitive functions may modify the clinical presentation of ADHD (Rommelse *et al.*, 2016), moderate its prognosis (Cheung *et al.*, 2015) and influence its clinical heterogeneity (Fair *et al.*, 2012; Mostert *et al.*, 2015), we included FIQ alongside the two main symptom dimensions of ADHD as the CCA symptom measures. FIQ is advantageous because it represents a parsimonious account of general cognitive functions, and is easily accessible in clinical routines, which was not the case for cognitive measures adopted in earlier attempts of clustering ADHD (Marquand *et al.*, 2016). Statistical significance at FWE-corrected alpha <0.05 was estimated via 100,000 permutations of the rows of one matrix relative to the other. Technical details regarding the CCA are provided in Supplementary Methods.

To test the existence of ADHD categorical biotypes, we implemented complementary analysis strategies using the features derived from CCA modes and combined features from connectivity and clinical symptoms. To assess whether the brain-symptom associations identified by the CCA were evenly distributed or clustered in distinct subgroups, we first employed *k*-means clustering on those linearly projected two-dimensional features by the CCA. To further test the existence of categorical biotypes, we explored joint clustering of connectivity features and symptom features using multi-view spectral clustering. This clustering algorithm maximizes the agreement across clusters identified in different feature spaces (connectivity and symptom) (Kumar and Daumé, 2011). The existence of valid discrete clusters detected using the *k*-means method was determined using state-of-the-art robustness measures including average silhouette width values and the Jaccard similarity (Hennig *et al.*, 2015), as well as the gap statistic (Tibshirani *et al.*, 2001). For the multi-view spectral clustering, we considered the convergence of the number of clusters yielded across different similarity thresholds. While there are no clear indications regarding the minimum sample size necessary for clustering analyses, broadly accepted guides (Hennig *et al.*, 2015) argue that our sample size (N=80) is substantially larger than that required to identify clusters using the number of features we employed. Methodological details regarding clustering algorithms are described in the Supplementary Methods.

## Results

### Altered whole-brain resting-state connectivity in medication-naive adults with ADHD

The application of NBS identified one significant brain network that differentiated adults with ADHD from healthy controls (FWE-corrected *p*=0.037; Fig. 2 and Supplementary Table 2&3 for details). The ADHD cohort showed stronger resting-state connectivity in this network compared to controls, with increased correlations between the DMN and frontoparietal network, the DMN and attention networks (including both salience/cingulo-opercular and dorsal attention components), the DMN and subcortical regions, the salience/cingulo-opercular network and sensory-motor and visual network, as well as the salience/cingulo-opercular network and dorsal attention, alongside frontoparietal networks) (Fig. 2). Confirmatory analyses using different brain parcellations yielded similar results (Supplementary Fig. 1).

**Figure 2.**
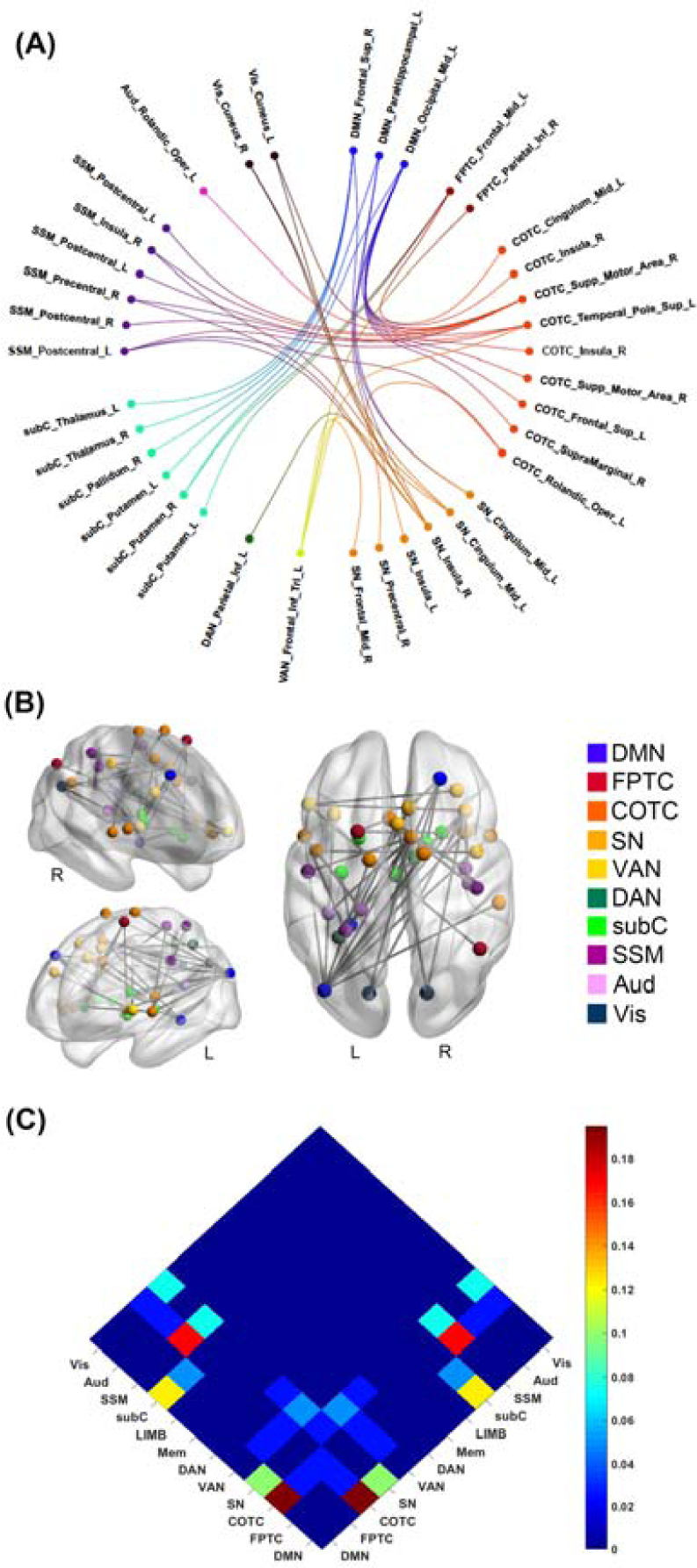
Group differences in inter-regional functional connectivity. The network-based statistic (NBS) identified a single network differentiating adult ADHD from healthy controls. This network was hyper-connected (i.e., higher positive correlations) in ADHD compared to controls. Functional brain networks were designated according to Power and colleagues (Power *et al.*, 2011). (A) Circular layout of the network distribution. Colors are used to define different brain networks. (B) Topological representation of brain network hyper-connectivity in ADHD compared to healthy controls. (C) The proportion of functional connections affected by ADHD. The number of connections involving each network pair was normalized by a total number of altered pair-wise connections. Adults with ADHD showed the highest positive functional connectivity between (i) the default mode (DMN) and the salience (SN) networks; (ii) the frontoparietal task control network (FPTC) and subcortical (subC) networks. Network nodes and edges (connections) were visualized using BrainNet Viewer (http://www.nitrc.org/projects/bnv/). Circular layouts were generalized with NeuroMArVL (http://immersive.erc.monash.edu.au/neuromarvl/). Networks/modules were assigned based on the community information provided by Power *et al.* (2011). The anatomical location was identified by the Automatic Anatomical Labeling atlas (http://www.gin.cnrs.fr/en/tools/aal-aal2/) and visual inspection. COTC=cingulo-opercular network; VAN=ventral attention network; DAN=dorsal attention network; SSM=somatosensorimotor network; Aud=auditory network; Vis=visual network; Supp=supplementary; R=right; L=left; Sup=superior; Inf=inferior; Mid=middle; Oper=opercular.

In a confirmatory analysis, we repeated the NBS analysis adjusting for subject-specific mean framewise displacements, gender, and age. In line with the original results, this analysis identified one significant brain network showing hyperconnectivity in adults with ADHD compared to healthy controls (Supplementary Fig. 2). This network was similar to the one detected by our main analysis (Fig. 2).

To test the stability of connectivity differences detected in our analysis, we undertook confirmatory analyses using different search thresholds (T values=3; 3.3; 3.9) for the NBS. As shown in Supplementary Fig. 3, the confirmatory NBSs identified similar patterns of hyperconnectivity between (i) the DMN and attention and cognitive control networks, (ii) the DMN and subcortical regions, and (iii) the salience/cingulo-opercular network and sensory processing networks in ADHD compared to healthy controls. Consistent with the effect of increasing the network forming threshold, these subnetworks become sparser as the serach threshold is increased.

The main NBS results were replicated using an alternative approach to characterize functional brain networks, using maps derived from ICA decomposition of the data rather than a predefined parcellation (Jafri *et al.*, 2008). Specifically, results from this additional analysis confirmed that ADHD had stronger connectivity between the DMN and the cingulo-opercular/salience/ventral attention networks compared to the controls (false discovery rate corrected *q*=0.044; Supplementary Fig. 4).

### ADHD biotyping

#### Dimensional biotype

The application of CCA identified one significant mode (*r*=0.430, FWE-corrected *p*=0.037) of interdependences between functional connectivity patterns and the symptom indices of hyperactivity-impulsivity, inattention, and general cognitive functioning (FIQ) (Supplementary Table 4a). Specifically, ADHD adults with high symptoms of hyperactivity and impulsivity showed higher resting-state connectivity in a network comprising the strongest increases in positive connectivity in ADHD compared to controls (Fig. 3A). Symptoms of inattention were only mildly positively associated with ADHD hyper-connectivity. On the other hand, general cognitive functions, as indexed by IQ scores, were inversely correlated with greater resting-state connectivity (Fig. 3A). The functional connections expressing the strongest positive associations in this mode (mean *r*=0.59, standard deviation=0.09) involved mainly altered DMN-cingulo-opercular and DMN-subcortical connectivity (Fig. 3B&C and Supplementary Table 5). The additional 2 modes were associated with very small effect sizes (r=0.18, r=0.02, respectively), and did not capture any additional meaningful association between patterns of brain connectivity and symptoms.

To test whether the CCA was not confounded by head motion, age or gender, we repeated this analysis using the significant subnetwork identified with the NBS in which these confounds were included as nuisance factors. This analysis also yielded one significant mode (r=0.415, FWE-corrected p=0.034; Supplementary Fig. 6), recapitulating the brain-symptom associations detected in the original analysis (Fig. 3).

**Figure 3.**
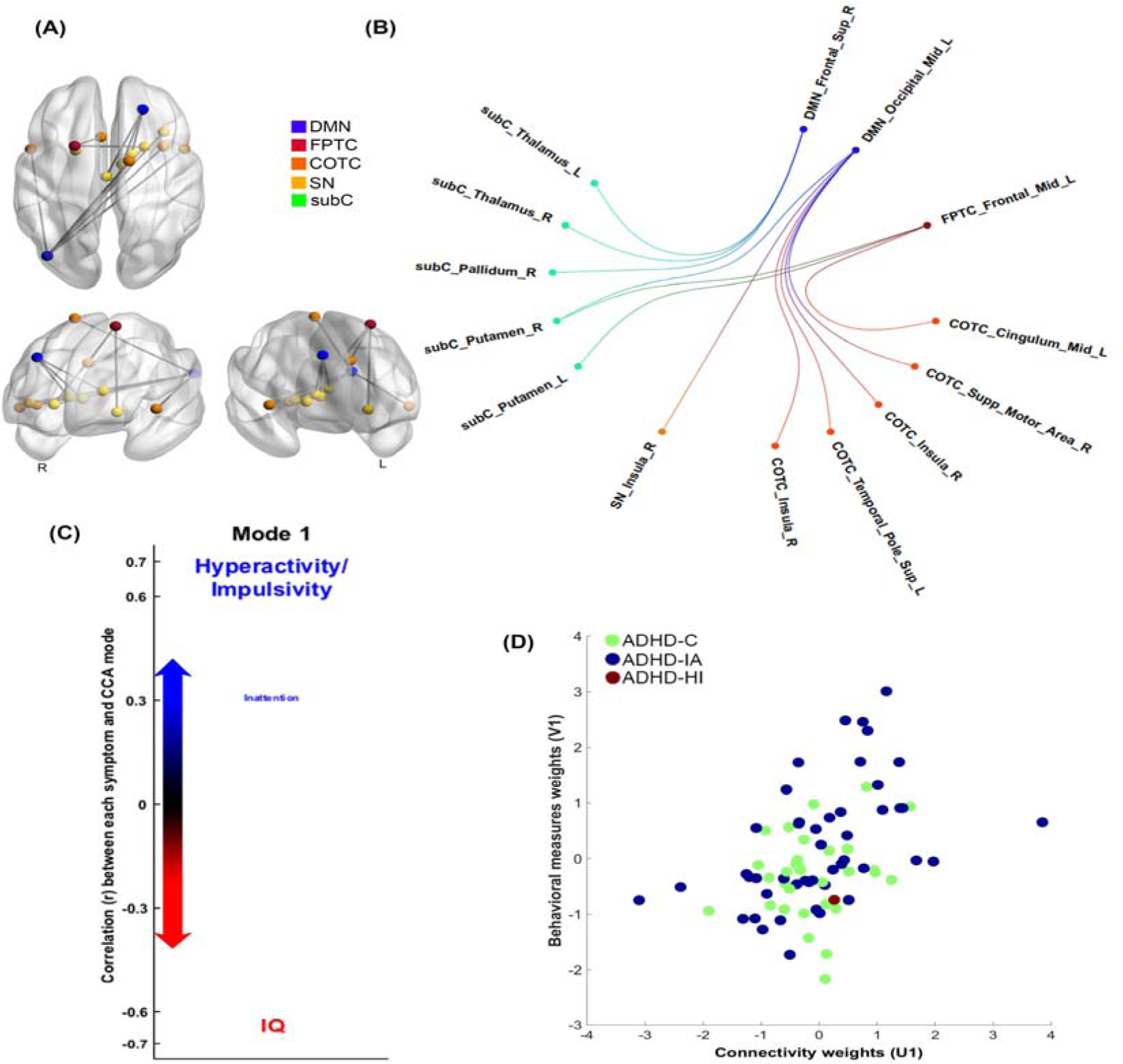
Canonical correlation analysis (CCA) and ADHD dimensional biotypes. (A) The CCA analysis identified a single significant (FWE-corrected p=0.037) mode of associations between resting-state connectivity and the behavioral variables of interest. The strength and direction of the variance explained by the CCA mode are indicated in the figure by the vertical position and font size. Adults with ADHD having predominant symptoms of hyperactivity and impulsivity showed higher resting-state connectivity in a network comprising the most (top 25%) hyper-connected edges in ADHD compared to controls (panel B). Conversely, higher cognitive functions (indexed by IQ scores) inversely correlated with (hyper) connectivity in the altered network. Symptoms of inattention were mildly positively associated with resting-state connectivity. (B) Representation of the top 25% node pairs expressed by the CCA mode. (C) This panel represents the circular layout of the top 25% node-pairs. This representation highlights that the majority of hyper-connected edges comprise functional interactions between the default mode (DMN) and the cingulo-opercular network (COTC) and subcortical (subC) networks. (D) Participants with different clinical ADHD subtypes were distributed evenly along the one-dimensional axis identified by the CCA. SN=salience network; FPTC=frontoparietal task control network; R=right; L=left; Sup=superior; Mid=middle; Supp=supplementary.

#### Categorical biotype

Results of clustering methods failed to show the existence of a non-overlapping ADHD categorical biotype. Specifically, the application of *k*-means clustering analysis to the CCA weighted symptom and connectivity values favored a dimensional solution over a categorical one. In particular, the value of the clustering index (average silhouette values) was 0.46 for 2 clusters and 0.49 for 3 clusters (Supplementary Table 6), both below the range considered to support a stable cluster solution. Based on the gap statistic (Supplementary Fig. 7), the suggested optimal number of clusters from the *k*-means algorithm was one. As shown in Supplementary Fig. 8A, ADHD participants were evenly distributed without any clear cluster demarcation. In support of this observation, a multi-view spectral clustering analysis using combined symptom and connectivity features also failed to converge to a stable clustering solution (Supplementary Fig. 8B). The inconsistency in the number of clusters detected across different similarity thresholds indicates that no valid categorical biotypes could be identified for adult ADHD.

### Comparison between DSM-IV subtypes and ADHD biotype

NBS and functional network connectivity analyses did not reveal significant differences in resting-state connectivity between the combined (N=32) and inattentive (N=47) clinical subtypes of ADHD.

We also assessed how the DSM-IV-defined ADHD clinical subtypes mapped on the brain-symptom dimensional axis revealed by our biotyping analysis. As shown in Fig. 3D, participants with different clinical ADHD subtypes were distributed evenly along the one-dimensional axis identified by the CCA.

### Gender effects on ADHD biotype

NBS did not identify significant connectivity differences between males and females with ADHD. As shown in Supplementary Fig. 9, males and females, regardless the clinical subtypes, were also distributed evenly along the one-dimensional axis identified by the CCA.

## Discussion

ADHD is characterized by substantial clinical and cognitive heterogeneity. Here, we questioned whether this childhood-onset adult ADHD heterogeneity could be parsed into dimensional or categorical biotypes. Our multivariate analyses showed that inter-individual differences in brain connectivity, clinical symptoms and general cognitive functioning in ADHD define a dimensional biotype (Fig. 3A). We found no evidence for a categorical definition of adult ADHD. Results from this study support the notion of adult ADHD as a single pathological entity, with altered brain-symptom associations varying across a single dimension or spectrum.

Data from our sample allowed the isolation of ADHD biotypes, which are not confounded by developmental delays, general cognitive dysfunction, history of medication use or confounds related to multi-site experiments. Using state-of-the-art multi-echo rs-fMRI data, we showed reduced functional segregation (i.e., higher positive functional connectivity) between the DMN and attention, alongside cognitive control networks in ADHD compared to healthy controls. One of the most robust findings of the fMRI literature in children and adults with ADHD is the increased functional interplay between the DMN and other networks of the brain, especially the cingulo-opercular/salience/ventral attention networks (Castellanos and Aoki, 2016; Gallo and Posner, 2016). In healthy controls, these between-network interactions are known to support dynamic switches between internal and external mental processes (Menon, 2011). Accordingly, studies have shown that the reduced functional segregation between the DMN and other brain networks correlated with attention and hyperactivity symptoms of ADHD (Cocchi *et al.*, 2012). For example, reduced functional segregation between the DMN and cognitive control networks has been linked to inattention (Barber *et al.*, 2015) and poor inhibitory control (van Rooij *et al.*, 2015) in ADHD youths. Also, the pharmacological enhancement of this between-networks functional segregation improves ADHD symptoms (Lin and Gau, 2015). The current study extends these previous findings to a larger sample of medication-naïve adults with clinical ADHD. Both NBS and functional network connectivity analyses show increased interactions between DMN and attention alongside cognitive control networks (especially the salience/cingulo-opercular component), supporting the notion that this deficit represents a key brain signature of ADHD (Castellanos and Aoki, 2016; Cai *et al.*, 2017). We also identified increased connectivity between salience/cingulo-opercular and all other major brain networks, as well as increased connectivity within the salience/cingulo-opercular network in adults with ADHD. This result supports the notion that altered salience network function significantly impacts the balance between brain systems activity supporting external and internal mental processes (Menon, 2011), and contributes to the emergence of ADHD symptoms (Castellanos and Aoki, 2016; Cai *et al.*, 2017).

The CCA showed one significant mode of the population variation that links a pattern of brain connectivity to a specific pattern of covariance between core ADHD symptoms and general intelligence. This analysis suggests that the heterogeneity of ADHD symptomatology can be described by a dimensional biotype. Symptoms of hyperactivity-impulsivity and inattention loaded onto positive associations with connectivity patterns in this mode, whereas IQ loaded onto negative associations. This result, despite being correlational in nature, echoes the earlier findings suggesting that preserved cognitive function may act as a protective factor for ADHD symptoms in adults (Cheung *et al.*, 2015; Rommelse *et al.*, 2016), whereas lower IQ is associated with chronic and persistent symptoms (Cheung *et al.*, 2015; Keyes *et al.*, 2017). Longitudinal studies would be required to better tease out any causal role in these associations. The result that hyperactivity-impulsivity and IQ loaded onto opposing ends of the same axis is in line with previous data on adolescence (Keyes *et al.*, 2017) and highlights the presence of this relationship in adulthood. Moreover, the fact that hyperactivity-impulsivity and inattention loaded onto the same CCA axis supports existing work suggesting a mildly positive correlation between these two ADHD symptom dimensions (Willcutt *et al.*, 2012). The functional connectivity patterns that were most strongly expressed by this mode included connections between DMN and cingulo-opercular and subcortical brain regions, as well as fronto-parietal-subcortical regions. This further supports proposals derived from healthy cohorts that anti-correlated activity between the DMN and cognitive-control networks underpins optimal brain functions and behaviors (Menon, 2011). Together, results from our CCA analysis highlight that a dimensional biotype defined by symptom-connectivity co-variation explains ADHD heterogeneity.

We revisited prior efforts to utilize neuroimaging or behavioral information to assess the existence of distinct subtypes of individuals with ADHD. Existing studies have defined several ADHD subtypes, mostly in the child population, characterized by co-occurring psychiatric symptoms (Acosta *et al.*, 2008), neuropsychological profiles (Fair *et al.*, 2012; Mostert *et al,* 2015), temperamental features (Karalunas *et al.*, 2014), and functional connectivity patterns (Gates *et al.*, 2014; Costa Dias *et al.*, 2015). However, these studies have failed to provide a consistent non-overlapping parcellation of ADHD.

Herein, we attempted to identify categorical ADHD biotypes by combining resting-state functional connectivity and symptom data in the largest sample of childhood-onset adult ADHD to date, but found no evidence for categorical biotypes. In keeping with the inconclusive results of previous attempts, our work highlights the difficulty in parsing the clinical heterogeneity of ADHD into non-overlapping categories. Whereas it could be argued that our clinically homogenous sample may have precluded the identification of categorical biotypes, the distribution of the current sample is consistent with demographic features of adults with childhood-onset ADHD (Asherson *et al.*, 2016). Likewise, our sample allowed the identification of ADHD-specific brain alterations and biotypes. Future studies may build upon the present results to interrogate the impact of factors such as comorbidity, developmental differences, and medication on the definition of ADHD biotypes. These multi-site endeavors will necessarily require significantly larger samples compared to the present work (Schnack and Kahn, 2016). Besides sample characteristics, we acknowledge that our results are linked to the symptoms included, and that we cannot exclude the possibility that ADHD heterogeneity could be parsed using different measures. It is therefore possible that more specific cognitive measures, such as those adopted in previous attempts (Fair *et al.*, 2012; Mostert *et al.*, 2015), may support the co-existence of categorical biotypes.

Intriguingly, we did not identify functional connectivity differences between the DSM-IV subtypes. Moreover, individuals across both clinically defined ADHD subtypes (i.e., combined and inattentive) were distributed evenly along the axis identified by the CCA, highlighting the challenges of clustering ADHD in non-overlapping subgroups (Marquand *et al.*, 2016). Whether our findings could be generalized to recently-identified late-onset ADHD (Faraone and Biederman, 2016) or children and adolescents with ADHD awaits further testing.

In the current study, we acquired multi-echo rs-fMRI, which allows direct measurement of T2* relaxation rates to facilitate the disambiguation of brain signals from noise and artifacts based on their different echo time dependence (Kundu *et al.*, 2012). The adopted analysis pipeline (see Supplementary Methods) has proven effective in denoising fMRI signal from motion and physiological artifacts in task and rest conditions (Kundu *et al.*, 2017). Despite advantages of our approach in detecting neural signals, we acknowledge the trade-off in acquiring multi-echo data by sacrificing some levels of spatial and temporal resolutions. Future studies are required to assess if dynamic functional connectivity could provide new insights into the link between brain and behavior in ADHD. For CCA, it is also important to note that, due to its correlational nature, CCA does not allow causal relationships to be inferred between brain and behavioral variables. Moreover, classic CCA, as implemented here, rests upon the assumptions of normality, which is the case herein. Where such assumptions are violated, variants such as sparse CCA, may be more suited.

Several limitations should be considered in addition to those considered above. The study sample was recruited only from one medical center in Taiwan and excluded patients with comorbid psychiatric conditions and psychotropic exposures. Future studies will need to assess the generalizability of our results to patients with such comorbidities as these are not uncommon in clinical practice. Second, only one participant presented with the hyperactive-impulsive subtype of ADHD. This ratio was consistent with the observation that the hyperactive-impulsive subtype is infrequent in adult ADHD (Asherson *et al.*, 2016). Whether this subtype of ADHD fits within the dimensional biotype identified herein would require further testing using a sample skewed to include more patients with this symptom profile. Lastly, although the gender ratio of the current adult ADHD cohort was typical (Asherson *et al.*, 2016), the small number of females might have precluded the unequivocal assessment of putative gender effects on biotyping.

In summary, we here provide the first biotyping of medication-naïve adults with ADHD using neuroimaging and symptom data. Results showed that patterns of co-variation between resting-state functional connectivity and clinically tractable symptom measures define a dimensional biotype. Despite the heterogeneity of its clinical presentation, the present work supports the notion of childhood-onset adult ADHD as a unitary disorder. Specifically, our findings highlight the need to consider its dimensional mediators in research and clinical interventions. This view is endorsed by a recent genome-wide association meta-analysis (Demontis *et al.*, 2017). As a whole, our findings support the importance of mapping the link between brain-behavioral phenotypes and clinical diagnosis, echoing the Research Domain Criteria framework to revise clinically defined constructs using a neurobiologically informed dimensional approach (Cuthbert, 2015).

## Acknowledgments

This work was supported by the Ministry of Technology and Science, Taiwan (MOST103-2314-B-002-021-MY3), the National Health Research Institutes, Taiwan (NHRI-EX103-10008PI), and National Taiwan University Hospital (NTUH103-S2458, NTUH104-S2761). The overseas fellowship of H.-Y.L. is supported by the Ministry of Technology and Science, Taiwan (106-2918-I-002-019), and National Taiwan University Hospital, Taipei, Taiwan. L.C. is supported by the Australian National Health Medical Research Council (L.C., APP1099082).

## Ethical Standards

The authors assert that all procedures contributing to this work comply with the ethical standards of the relevant national and institutional committees on human experimentation and with the Helsinki Declaration of 1975, as revised in 2008.

## 1. Supplementary Methods

### 1.1. Data

#### 1.1.1. Measures for ADHD symptoms

##### 1.1.1.1. The Adult ADHD Self-Report Scales

The Adult ADHD Self-Report Scales (ASRS), an 18-question scale, was developed in conjunction with the revision of the World Health Organization (WHO) Composite International Diagnostic Interview (CIDI). The ASRS consists of two subscales, Inattention (nine items) and Hyperactivity-Impulsivity (nine items), according to the 18 *DSM-IV* ADHD symptom criteria. Each item asks how often a symptom occurred during the last 6 months on a 5-point Likert scale: 0=never, *1=rarely*, 2=*sometimes*, 3=*often*, and 4=very *often.* The psychometric properties of the Chinese ASRS have been established in a sample of 4,329 Taiwanese young adults (Yeh *et al.*, 2008). The intraclass correlations (ICCs) for test-retest reliability ranged from 0.80 for the Inattention subscale, 0.82 for the Hyperactivity-Impulsivity subscale, and .85 for the total score. The internal consistency (Cronbach’s α) was high for the Inattention subscale (0.87), the Hyperactivity-Impulsivity subscale (0.85), and the total score (0.91). It has been used in studies on adult ADHD and sleep problems, anxiety/depression symptoms, and quality of life in Taiwan (Gau *et al.*, 2007; Chao *et al.*, 2008).

##### 1.1.1.2. The Swanson, Nolan, and Pelham, Version IV Scale (SNAP-IV)-Parent form

The SNAP-IV is a 26-item rating instrument including the core DSM-IV-derived ADHD subscales of IA, HI and OD subscales (items 1-9, 10-18, and 19-26, respectively) (Swanson *et al.*, 2001). Each item is rated on a 4-point Likert scale, 0-3 for “not at all”, “just a little”, “quite a lot”, and “very much” based on and parents’ report. The norm and psychometric properties of the Chinese version of SNAP-IV have been well established in Taiwan by Gau and colleagues (Gau *et al.*, 2008). The scale has good test-retest reliability (ICCs 0.59~0.72), high internal consistency (Cronbach’s α>0.88) and discriminative validity (Gau *et al.*, 2008) and is commonly used in clinical evaluation and research in Taiwanese child and adolescent populations (Yang *et al.*, 2013).

##### 1.1.1.3. The modified adult version of the ADHD supplement of the Chinese version of the Schedule for Affective Disorders and Schizophrenia-Epidemiological Version (K-SADS-E)

The K-SADS-E is a semi-structured interview scale for the systematic assessment of both past and current episodes of mental disorders in children and adolescents (Orvaschel *et al.*, 1982). Development of the Chinese K-SADS-E was completed by the Child Psychiatry Research Group in Taiwan (Gau and Soong, 1999). This included a two-stage translation and modification for several items with psycholinguistic equivalents relevant to the Taiwanese culture and further modification to meet the DSM-IV diagnostic criteria, with high reliability (generalized kappa coefficients ranging from 0.73 to 0.96 for all mental disorders) and validity (sensitivity 78% and specificity 98%) (Gau *et al.*, 2005). In order to obtain the information about ADHD symptoms and diagnoses in adulthood according to the DSM-IV diagnostic criteria, semi-structured interviews were conducted using both the modified adult ADHD supplement and the Conners’ Adult ADHD Diagnostic Interview for DSM-IV (Takahashi *et al.*, 2014). The results showed that the ADHD diagnosis in childhood and current adulthood based on the two clinical instruments achieved total agreement (i.e., people who had been diagnosed with ADHD in childhood and/or current adulthood using the modified adult ADHD supplement of the K-SADS-E also acquired the ADHD diagnosis based on the Conners’ Adult ADHD Diagnostic Interview).

### 1.2. Analyses

#### 1.2.1. Multi-echo independent component analysis (ME-ICA)

ME-ICA initially decomposed multi-echo rs-fMRI data into independent components using FastICA (Hyvarinen, 1999). Independent components were subsequently categorized as BOLD or non-BOLD components based on Kappa and Rho values, which were yielded from signal models reflecting the BOLD-like or non-BOLD-like signal decay processes (Kundu *et al.*, 2012). BOLD-related signals show linear dependence of percent signal changes on TE, which is the characteristic of the T2* decay (Huettel *et al.*, 2008). On the other hand, non-BOLD signal amplitudes demonstrate TE-independence. TE dependence of BOLD signal was measured using the pseudo-F-statistic Kappa, with components that scaled strongly with TE having high Kappa scores. Non-BOLD components were identified by TE independence measured by the pseudo-F-statistic Rho. By removing non-BOLD components, data were denoised for head motion, physiological, and scanner artifacts (Kundu *et al.*, 2013).

#### 1.2.2. Functional network connectivity analysis

##### 1.2.2.1. Independent Component Analysis (ICA)

After preprocessing, the temporally concatenated probabilistic ICA algorithm (temporally concatenated) implemented in FSL MELODIC (Beckmann and Smith, 2004) was used to analyze the rs-fMRI data of all participants. Non-brain voxels were masked with voxel-wise demeaning of the data and normalization of the voxel-wise variance. Next, the processed data were whitened and projected into a 20-dimensional subspace using a Principal Components Analysis (PCA). This step provided a fine-grained decomposition of interconnected brain regions (Smith *et al.*, 2009). These whitened observations were decomposed into sets of vectors that describe signal variation across (i) the temporal domain (time courses), (ii) the session/subject domain, and (iii) the spatial domain (spatial maps). This decomposition was implemented through a non-Gaussian spatial source distribution using a fixed-point iteration technique (Hyvarinen, 1999). Estimated component maps were divided by the standard deviation of the residual noise, with a threshold of 0.5 set (the probability that needed to be exceeded by a voxel to be considered ‘active’ in the component of interest) by fitting a mixture model to the histogram of intensity values (Beckmann and Smith, 2004).

We selected resting-state networks according to their known spatial distribution (Smith *et al.*, 2009; Yeo *et al.*, 2011; Cocchi *et al.*, 2012). We extracted 20 ICA components, 14 of which are consistently identified as canonical resting-state networks (Yeo *et al.*, 2011). The similarity of these 14 resting-state networks with those previously identified was quantified using spatial correlation (all spatial correlation values >0.4) and confirmed by visual inspection. Only these 14 components (networks) were considered in subsequent analyses (Supplementary Fig. 2A).

##### 1.2.2.2. Functional Network Connectivity

The summary time-course for each resting-state network was calculated at the individual participant’s level by spatial regression of the full set of 20 ICA components against each participant’s denoised rs-fMRI data. This approach models are overlapping variance to account for the potential effects of residual noise captured by the non-physiological valid components (N=6). We calculated functional network connectivity (FNC) (Jafri *et al.*, 2008; Lv *et al.*, 2016) using the Pearson correlation coefficient between each other summary time course. This resulted in a 3D FNC matrix with the dimensions of 14 × 14 (networks) × 203 (participants). Group differences in FNC were tested for each pair of networks using one-way analysis of variance (ANOVA), and FNC with significant group differences were further tested by 2-sample t-test to determine the direction of the difference. The significance threshold was set at q<0.05, corrected for multiple comparisons using false discovery rate (FDR) (Benjamini and Hochberg, 1995).

### 1.2.3. Principal component analysis for ADHD symptoms

To circumvent reporting biases (Asherson *et al.*, 2016), core ADHD symptoms were encapsulated as factor scores (DiStefano *et al.*, 2009) derived from a principal component analysis (PCA) of self-, parents-, and clinician-reported measures, including self-rated Adult ASRS (Yeh *et al.*, 2008), parent-rated SNAP-IV (Gau *et al.*, 2008), as well as a clinician-rated modified adult version of the ADHD supplement of the Chinese version of the K-SADS-E (Chang *et al.*, 2013; Ni *et al.*, 2013a; Ni *et al.*, 2013b) (the number of ADHD measures used in the PCA was 3). Two principal components were extracted, which explained 88.25% of the total variance. Factors were orthogonalized using Varimax rotation. Among them, the first component explained 51.58% of the total variance, and all of the hyperactivity-impulsivity subscales from the above three measures were consistently loaded on this component. The second component explained 36.67% of the total variance, and scores of inattention subdomain across 3 measures were loaded on the component. The principal component analysis was implemented using IBM SPSS Statistics for Macintosh, Version 22.0 (IBM Corp., Armonk, NY, USA).

*Symptoms scores patterns loaded onto two components (rotated component matrix)*

|  | Component |  |
| --- | --- | --- |
|  | 1 (Hyperactivity-impulsivity) | 2 (Inattention) |
| SNAP_Inattention | 0.244 | 0.743 |
| SNAP_Hyperactivity-impulsivity | 0.842 | 0.182 |
| K-SADS-E_Inattention | 0.224 | 0.806 |
| K-SADS-E_Hyperactivity-impulsivity | 0.798 | 0.172 |
| ASRS-A | 0.227 | 0.871 |
| ASRS-B | 0.754 | 0.416 |

### 1.2.4. Canonical correlation analysis (CCA)

Both connectivity and behavioral measures were normalized and demeaned. A further regression of in-scanner head motion confounds also performed following the approach of Smith and colleagues (Smith *et al.*, 2015) (http://www.fmrib.ox.ac.uk/analysis/HCP-CCA). To avoid overfitting the CCA, a PCA was undertaken using the FSLNets toolbox (Smith *et al.*, 2014) to reduce the dimensionality of the deconfounded functional connectivity matrix to three eigenvectors (explaining 31.83% of the total variance in the connectivity matrix; Supplementary Fig. 5). The data was reduced to this resolution to keep the methodological steps as per Smith et al. (Smith *et al.*, 2015), given the three behavioral measures selected in the CCA. We note that no consensus exists for component number selection (Abdi and Williams, 2010). Thus, we also employed a confirmatory CCA analysis based on a larger dimensionality of 5 eigenvectors (explaining 43.9% of the variance in the connectivity matrix). The primary (3 eigenvectors, *r*=0.430, FWE-corrected *p*=0.037) and confirmatory CCA (*r*=0.446, FWE-corrected *p*=0.049; Supplementary Table 4b) yielded similar results. Thus, only results from the primary CCA are reported in the main text.

We next assessed which functional connections were most strongly expressed by variations in the original sets of connections captured by each CCA mode. CCA provides an output vector describing the extent (weight) to which a given individual’s connectivity pattern correlated with the CCA mode. We correlated this vector against the original connectivity matrix identified by the NBS analysis to obtain a vector mapping the relative weights and directional signs of the association between resting-state connectivity and the CCA mode (weighted feature vector). In line with what previously done, the strongest (top 25%) absolute values in this vector were retained to define the strongest associations between individual connectivity weights and behavioral measures (Smith *et al.*, 2015).

### 1.2.5. Clustering algorithms for categorically subtyping ADHD

To test the existence of ADHD categorical biotypes, we implemented several complementary analyses using the connectivity and clinical features derived from the significant CCA mode, and combined features from connectivity and clinical symptoms, respectively.

#### 1.2.5.1. k-means clustering algorithm based on brain-behavior features derived from the significant CCA mode.

To assess whether the brain-behavior associations identified by the CCA could be clustered into non-overlapping subgroups, we first used *k*-means clustering on the features linearly projected by the CCA. This standard clustering procedure uses individual brain-behavior associations to assign each participant to exactly one of *k* clusters (based on clinical ADHD subtypes, a *k*=2 or 3 was used here) (Venkataraman *et al.*, 2009). To reach stable clustering results, for each setting of *k*, clustering was repeated for 10,000 times so that the participants-to-centroid distances within-cluster sum-of-squares were minimized.

#### 1.2.5.2. Multi-view spectral clustering algorithm based on features of functional connectivity and clinical symptoms

With regards to multi-view spectral clustering algorithm (Shi and Malik, 2000; Kumar and Daumé,2011), we considered clusters derived from the analysis of altered functional connectivity in ADHD compared to controls and features related to clinical symptoms/IQ as two views contributing to the clustering. Using the multi-view spectral clustering framework, the substantial variability of categorical subgrouping across multimodal features (connectivity and behavior) could be modeled and accounted for. This novel clustering method has the advantage of effectively addressing heterogeneity in the considered features by maximizing the agreement across multimodal clusters (Shi and Malik, 2000; Kumar and Daumé,2011).

Spectral clustering uses connectivity (*denoised NBS results*) and clinical features (*inattention, hyperactivity-impulsivity, and IQ*), respectively, to generate two graphs. Nodes within the graphs represent individuals with ADHD whereas the edges represent the similarities between nodes (individuals). The two graphs (one mapping connectivity and one mapping behavior) were then partitioned using the normalized cut strategy, in which the top *k* eigenvectors of the normalized graph Laplacian, which carries the most discriminative information, are adopted to cut the graphs into clusters efficiently. Subsequent co-training algorithms search for target clusters that predict same labels for co-occurring patterns in each view. The spectral clustering algorithm of bi-partitioning sub-graph stopped when the normalized cut value (representing the similarity between the subjects within each possible cluster) is larger than the pre-set threshold. There is no consensus regarding the optimal threshold to be used. Thus, we examined thresholds ranging from 0.2 to 0.9 (incremental of 0.1) (Chen *et al.*, 2013). We used 10,000 iterations for co-training algorithms to converge on stable clusters (permuting for each threshold).

#### 1.2.5.3. Validity of k-means clustering

We verify the validity of *k*-means clustering using average silhouette width values (Kononenko and Kukar, 2007), the Jaccard similarity (Hennig, 2008), and the gap statistic (Tibshirani *et al.*, 2001). This information is provided in Supplementary Table 6 (average silhouette width values and the Jaccard similarity) and Supplementary Fig. 7 (the Gap statistic).

The silhouette width value is a combination measure assessing intra-cluster homogeneity and inter-cluster separation. It is calculated by measuring how similar that point is to points in its own cluster when compared to points in other clusters. The cutoffs to interpret the validity of *k*-means clustering based on average silhouette width values are as follows (Kononenko and Kukar, 2007):

|  |  |
| --- | --- |
| 0.71-1.0 | A strong structure has been found. |
| 0.51-0.70 | A reasonable structure has been found. |
| 0.26-0.50 | The structure is weak and could be artificial. Try additional methods of data analysis. |
| <0.25 | No substantial structure has been found. |

Jaccard’s similarity (Hennig, 2008) is defined as the size of the intersection divided by the size of the union of the assigned clusters and the resulting partitions from resampling pipelines. It allows estimating the frequency with which similar clusters were recovered in the data. The clustering results with Jaccard’s similarity <0.5 are considered unstable (Hennig, 2008).

The gap statistic (Tibshirani *et al.*, 2001) standardizes the graph of log(*W_k_*), where *W_k_* is the within-cluster dispersion defined by the within-cluster sum of squares around the cluster means, by comparing it to its expectation under an appropriate null reference distribution of the data. The ‘k’ is the number of clusters. The estimate of the optimal number of clusters is defined by searching for the local maximum of the graph, and selecting the smallest *k* within one standard error of the local max.

#### 1.2.5.4. The issue of sample size for clustering analyses

There is no clear indication regarding the minimum sample size necessary for clustering analyses. However, it is suggested that the minimal sample size for clustering analyses should not be less than 2^m^ cases (*m*=number of features used), with 5*2^m^ considered preferable (Dolnicar, 2002). In the present study, we fed features linearly projected by the CCA (i.e., 1 for the brain connectivity feature; 1 for the behavior feature) into *k*-means clustering. That is, the minimum sample size for *k*-means clustering is 20 subjects (i.e., 5*2^2^=20). Concerning the multi-view spectral clustering, there has been very limited prior work investigating the minimum sample size required to obtain meaningful clusters. The multi-view spectral clustering algorithm is, however, considered robust for the high dimensionality and small-sample-size problem (Tao *et al.*, 2014). Indeed, a smaller sample size is generally required to obtain a solution (i.e., the most robust clustering results) using multi-view spectral clustering compared to single-view clustering (Kumar and Daumé, 2011).

In keeping with the above, the current sample size (N=80 ADHD) is appropriate for both *k*-means and spectral clustering (Dolnicar, 2002; Kumar and Daumé, 2011; Tao *et al.*, 2014).

## 2. Supplementary Tables

**Supplementary Table 1a.** Demographics among attention-deficit hyperactivity subtypes (ADHD) (based on the current presentation of ADHD psychopathology)

| Mean (SD) | ADHD-C (N=32) | ADHD-I (N=47) | ADHD-H (N=1) | Statistics<sup>c</sup> |
| --- | --- | --- | --- | --- |
| Age | 27.5 (5.2) | 26.3 (5.9) | 19.7 | $p=0.352$ |
| Sex (M/F) | 25/7 | 30/17 | 1/0 | $p=0.319$ |
| Handedness (R/L) | 24/8 | 38/9 | 1/0 | $p=0.800$ |
| FIQ | 108.8 (8.3) | 106.4 (11.5) | 115 | $p=0.280$ |
| VIQ | 108.8 (11.0) | 104.1 (11.1) | 103 | $p=0.069$ |
| PIQ | 110.3 (9.9) | 106.7 (19.1) | 128 | $p=0.320$ |
| *ADHD symptoms* |  |  |  |  |
| Inattention<sup>a</sup> | 21.4 (3.8) | 18.6 (5.4) | 3 | $p=0.011$ |
| Hyperactivity/Impulsivity<sup>a</sup> | 17.6 (6.4) | 11.0 (5.0) | 13 | $p<0.001$ |
| Opposition-defiance<sup>a</sup> | 13.8 (5.3) | 9.9 (5.7) | 7 | $p=0.003$ |
| ASRS-A | 28.5 (3.4) | 26.4 (4.9) | 4 | $p=0.037$ |
| ASRS-B | 24.2 (4.0) | 17.4 (5.5) | 12 | $p<0.001$ |
| Mean frame-wise displacement<sup>b</sup> (mm) | 0.050 (0.025) | 0.048 (0.025) | 0.049 | $p=0.737$ |

**Supplementary Table 1b.**
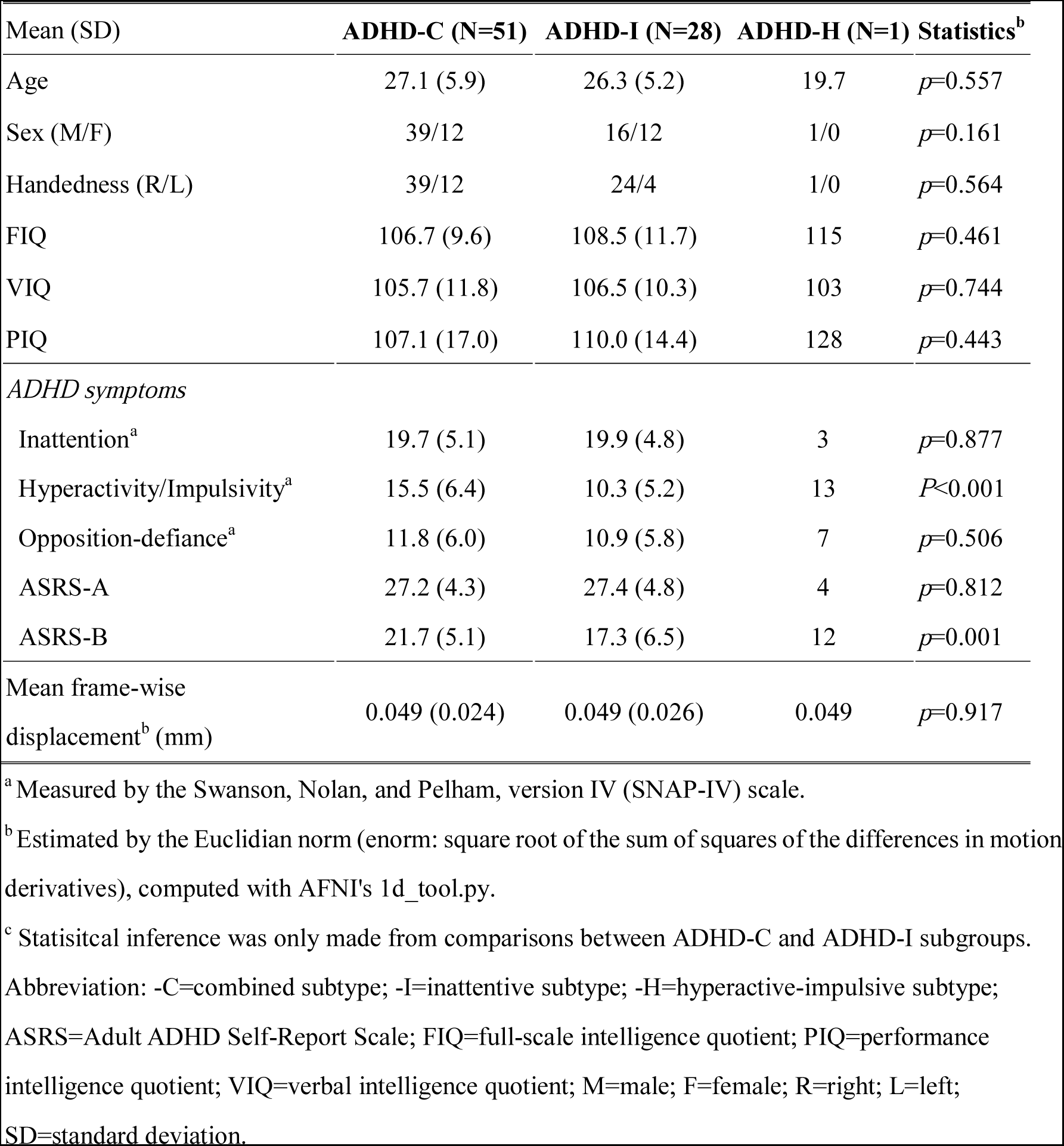
Demographics among attention-deficit hyperactivity (ADHD) subtypes (based on the childhood presentation of ADHD psychopathology)

| Mean (SD) | ADHD-C (N=51) | ADHD-I (N=28) | ADHD-H (N=1) | Statistics<sup>b</sup> |
| --- | --- | --- | --- | --- |
| Age | 27.1 (5.9) | 26.3 (5.2) | 19.7 | $p=0.557$ |
| Sex (M/F) | 39/12 | 16/12 | 1/0 | $p=0.161$ |
| Handedness (R/L) | 39/12 | 24/4 | 1/0 | $p=0.564$ |
| FIQ | 106.7 (9.6) | 108.5 (11.7) | 115 | $p=0.461$ |
| VIQ | 105.7 (11.8) | 106.5 (10.3) | 103 | $p=0.744$ |
| PIQ | 107.1 (17.0) | 110.0 (14.4) | 128 | $p=0.443$ |
| *ADHD symptoms* |  |  |  |  |
| Inattention<sup>a</sup> | 19.7 (5.1) | 19.9 (4.8) | 3 | $p=0.877$ |
| Hyperactivity/Impulsivity<sup>a</sup> | 15.5 (6.4) | 10.3 (5.2) | 13 | $P<0.001$ |
| Opposition-defiance<sup>a</sup> | 11.8 (6.0) | 10.9 (5.8) | 7 | $p=0.506$ |
| ASRS-A | 27.2 (4.3) | 27.4 (4.8) | 4 | $p=0.812$ |
| ASRS-B | 21.7 (5.1) | 17.3 (6.5) | 12 | $p=0.001$ |
| Mean frame-wise displacement<sup>b</sup> (mm) | 0.049 (0.024) | 0.049 (0.026) | 0.049 | $p=0.917$ |

<sup>a</sup> Measured by the Swanson, Nolan, and Pelham, version IV (SNAP-IV) scale.
<sup>b</sup> Estimated by the Euclidian norm (enorm: square root of the sum of squares of the differences in motion derivatives), computed with AFNI's 1d\_tool.py.
<sup>c</sup> Statistical inference was only made from comparisons between ADHD-C and ADHD-I subgroups.
Abbreviation: -C=combined subtype; -I=inattentive subtype; -H=hyperactive-impulsive subtype;
ASRS=Adult ADHD Self-Report Scale; FIQ=full-scale intelligence quotient; PIQ=performance
intelligence quotient; VIQ=verbal intelligence quotient; M=male; F=female; R=right; L=left;
SD=standard deviation.

**Supplementary Table 2.** Details of the nodes within the altered network of ADHD (network-based statistics, NBS)

|  | **MNI coordinates** |  |  |
| --- | --- | --- | --- |
| **Nodes** | x | y | z |
| DMN_Frontal_Sup_R | 22 | 39 | 39 |
| DMN_Occipital_Mid_L | -41 | -75 | 26 |
| DMN_ParaHippocampal_L | -26 | -40 | -8 |
| FPTC_Frontal_Mid_L | -23 | 11 | 64 |
| FPTC_Parietal_Inf_R | 44 | -53 | 47 |
| SN_Precentral_R | 42 | 0 | 47 |
| SN_Insula_L | -35 | 20 | 0 |
| SN_Insula_R | 36 | 22 | 3 |
| SN_Cingulum_Mid_L | -1 | 15 | 44 |
| SN_Frontal_Mid_R | 31 | 33 | 26 |
| SN_Cingulum_Mid_L | 5 | 23 | 37 |
| COTC_Frontal_Sup_L | -16 | -5 | 71 |
| COTC_SupraMarginal_R | 54 | -28 | 34 |
| COTC_Rolandic_Oper_L | -45 | 0 | 9 |
| COTC_Supp_Motor_Area_R | 13 | -1 | 70 |
| COTC_Insula_R | 49 | 8 | -1 |
| COTC_Temporal_Pole_Sup_L | -51 | 8 | -2 |
| COTC_Supp_Motor_Area_R | 7 | 8 | 51 |
| COTC_Insula_R | 36 | 10 | 1 |
| COTC_Cingulum_Mid_L | -5 | 18 | 34 |
| SN_Frontal_Mid_R | 31 | 33 | 26 |
| SN_Cingulum_Mid_L | 5 | 23 | 37 |
| COTC_Frontal_Sup_L | -16 | -5 | 71 |
| COTC_SupraMarginal_R | 54 | -28 | 34 |
| COTC_Rolandic_Oper_L | -45 | 0 | 9 |
| COTC_Supp_Motor_Area_R | 13 | -1 | 70 |
| COTC_Insula_R | 49 | 8 | -1 |
| COTC_Temporal_Pole_Sup_L | -51 | 8 | -2 |
| COTC_Supp_Motor_Area_R | 7 | 8 | 51 |
| COTC_Insula_R | 36 | 10 | 1 |
| COTC_Cingulum_Mid_L | -5 | 18 | 34 |
| DAN_Parietal_Inf_L | -33 | -46 | 47 |
| VAN_Frontal_Inf_Tri_L | -49 | 25 | -1 |
| subC_Putamen_L | -22 | 7 | -5 |
| subC_Putamen_R | 23 | 10 | 1 |
| subC_Pallidum_R | 15 | 5 | 7 |
| subC_Thalamus_R | 9 | -4 | 6 |
| subC_Thalamus_L | -2 | -13 | 12 |
| Vis_Cuneus_R | 15 | -77 | 31 |
| Vis_Cuneus_L | -16 | -77 | 34 |

Abbreviations: ADHD=attention-deficit hyperactivity disorder; MNI=Montreal Neurological Institute; DMN=default mode network; SN=salience network; COTC=cingulo-opercular network; FPTC=frontoparietal task control network; VAN=ventral attention network; DAN=dorsal attention network; SSM=somatosensorimotor network; Aud=auditory network; Vis=visual network; subC=subcortical; Supp=supplementary; R=right; L=left; Sup=superior; Inf=inferior; Mid=middle; Oper=opercular.

**Supplementary Table 3.**
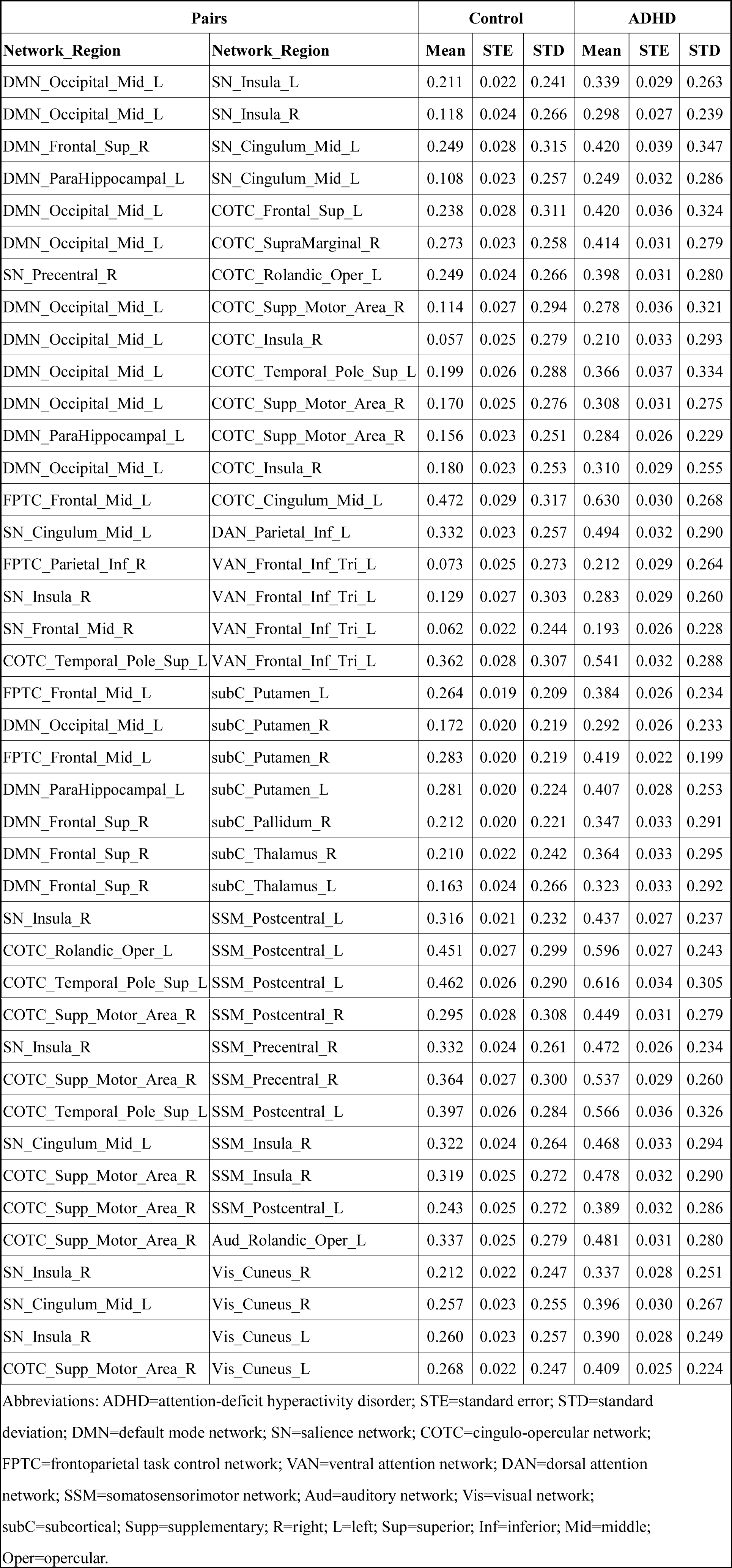
Average values of functional connectivity in the pairwise connections of interest (network-based statistics, NBS)

**Supplementary Table 4a.** The significant canonical correlation analysis (CCA) mode (*p*<0.05, family-wise error corrected) of the primary analysis.

|  |  |
| --- | --- |
| **CCA mode** | One |
| $df_1$ | 9 |
| $df_2$ | 180.25 |
| $F$ | 2.03 |
| $r$ | 0.430 |
| *Wilk's lambda* | 0.7740 |
| *Familywise error corrected p* | 0.0367 |

**Supplementary Table 4b.** The significant CCA mode (*p*<0.05, family-wise error corrected) based on the 5 eigenvectors derived from the connectivity matrix.

|  |  |
| --- | --- |
| **CCA mode** | One |
| $df_1$ | 15 |
| $df_2$ | 199.17 |
| $F$ | 1.61 |
| $r$ | 0.446 |
| *Wilk's lambda* | 0.7030 |
| *Familywise error corrected p* | 0.0491 |

**Supplementary Table 5.** Canonical correlation analysis (CCA) mode connectivity weight and associated interregional pairs

| Pairs |  | CCA edge strength modulation |
| --- | --- | --- |
| Network_Region | Network_Region |  |
| DMN_Occipital_Mid_L | SN_Insula_R | 0.549 |
| DMN_Occipital_Mid_L | COTC_Supp_Motor_Area_R | 0.656 |
| DMN_Occipital_Mid_L | COTC_Insula_R | 0.757 |
| DMN_Occipital_Mid_L | COTC_Temporal_Pole_Sup_L | 0.755 |
| DMN_Occipital_Mid_L | COTC_Insula_R | 0.623 |
| FPTC_Frontal_Mid_L | COTC_Cingulum_Mid_L | 0.568 |
| FPTC_Frontal_Mid_L | subC_Putamen_L | 0.510 |
| DMN_Occipital_Mid_L | subC_Putamen_R | 0.643 |
| FPTC_Frontal_Mid_L | subC_Putamen_R | 0.510 |
| DMN_Frontal_Sup_R | subC_Pallidum_R | 0.502 |
| DMN_Frontal_Sup_R | subC_Thalamus_R | 0.500 |
| DMN_Frontal_Sup_R | subC_Thalamus_L | 0.564 |

Abbreviations: DMN=default mode network; SN=salience network; COTC=cingulo-opercular network; FPTC=frontoparietal task control network; subC=subcortical; R=right; L=left; Sup=superior; Mid=middle; Supp=supplementary.

**Supplementary Table 6.** Validity indices of *k*-means clustering method (based on the feature vectors of individual participant’s weight derived from the connectivity and symptoms matrices of canonical correlation analysis)

| **$k$-means clustering** |  |  |
| --- | --- | --- |
|  | 2<br>clusters | 3<br>clusters |
| Jaccard similarity | 0.3385 | 0.5737 |
| Average silhouette width values | 0.4640 | 0.4861 |

## 3. Supplementary Figures

**Supplementary Figure 1.**
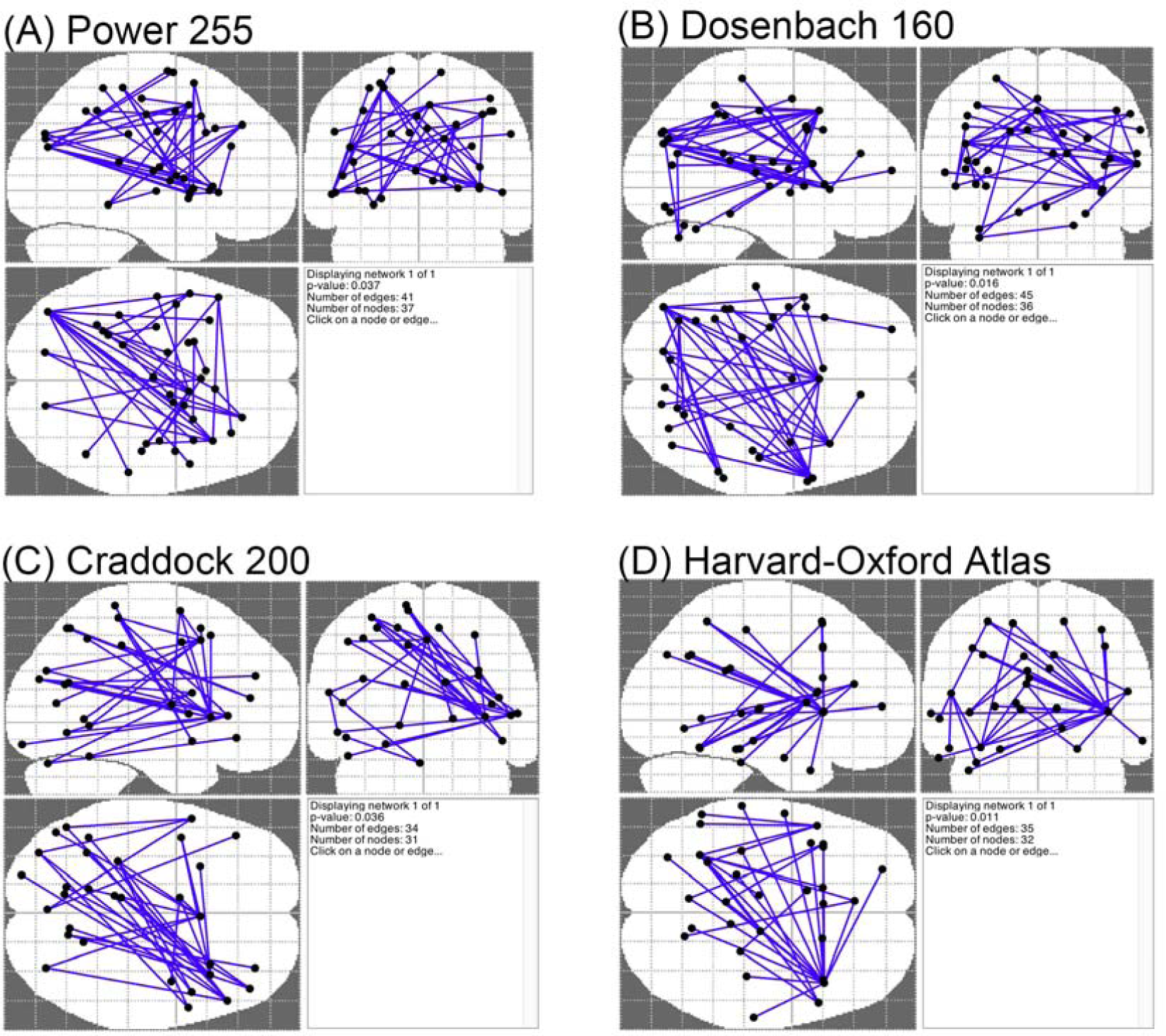
Changes in functional connectivity between adult ADHD and matched healthy controls across different brain parcellations. The network-based statistic (NBS) showed stronger (generally stronger positive correlations, see Supplementary Table 3) functional connectivity in a single whole-brain network in ADHD compared to healthy controls. (A) 255 regions of interest parcellation (Power *et al.*, 2011). (B) 160 regions of interest parcellation (Dosenbach *et al.*, 2010). (C) 200 regions of interest parcellation (Craddock *et al.*, 2012). (D) the anatomical parcellation based on The Harvard-Oxford probabilistic cortical and subcortical atlases (www.fmrib.ox.ac.uk/fsl). Overall, the results obtained from different brain parcellations were similar.

**Supplementary Figure 2.**
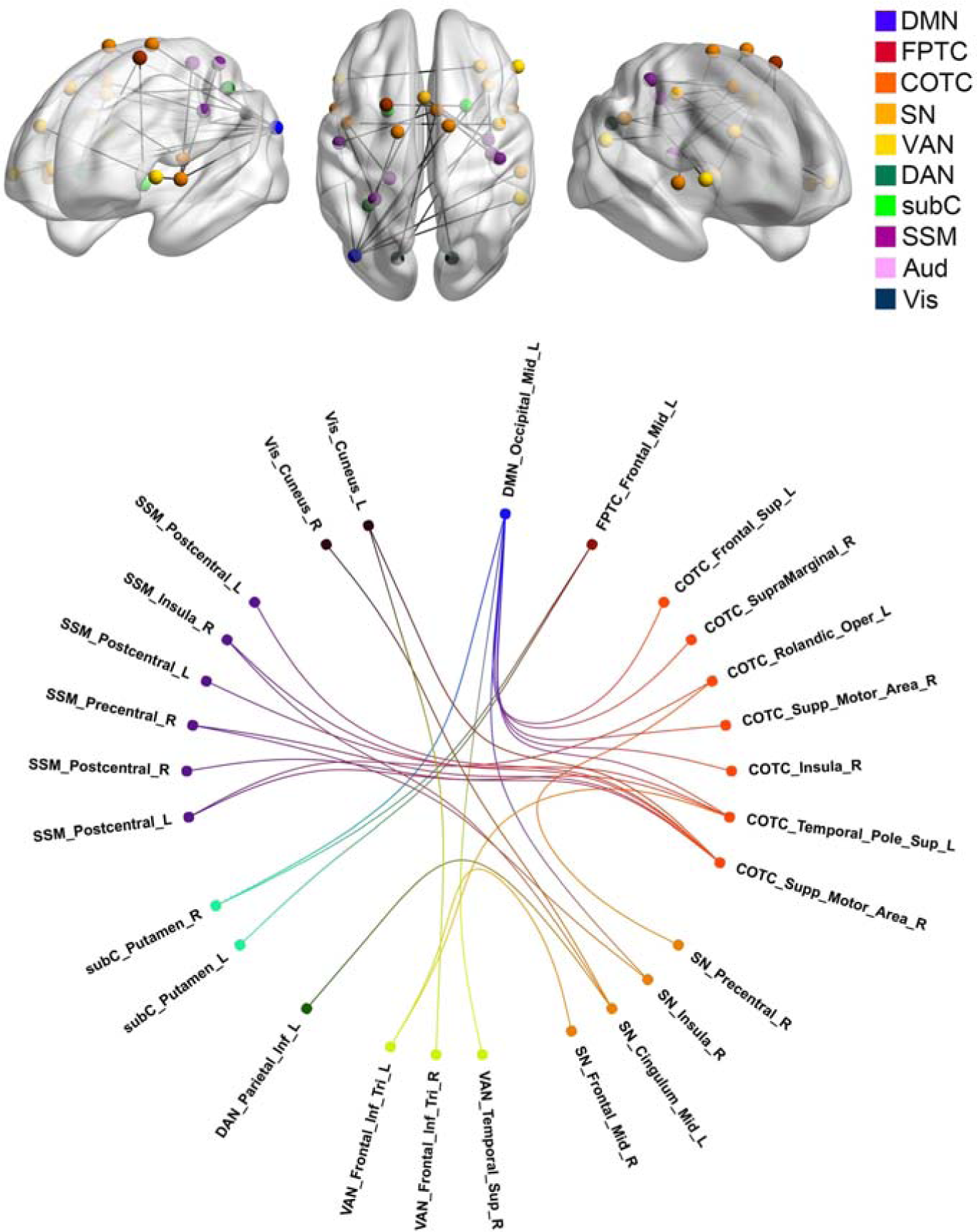
With additional adjusting for demographic features, group differences in inter-regional functional connectivity. The network-based statistic (NBS) adjusting for gender/sex, levels of in-scanner head motion, and age identified a single network differentiating adult ADHD from healthy controls. This network was of largely the same pattern with the main analysis as shown in Figure 2: Adults with ADHD showed increased correlations between the DMN and frontoparietal network, the DMN and attention networks (including both salience/cingulo-opercular and dorsal attention components), the DMN and subcortical regions, the salience/cingulo-opercular network and sensory-motor and visual network, as well as the salience/cingulo-opercular network and dorsal attention alongside frontoparietal networks.

**Supplementary Figure 3.**
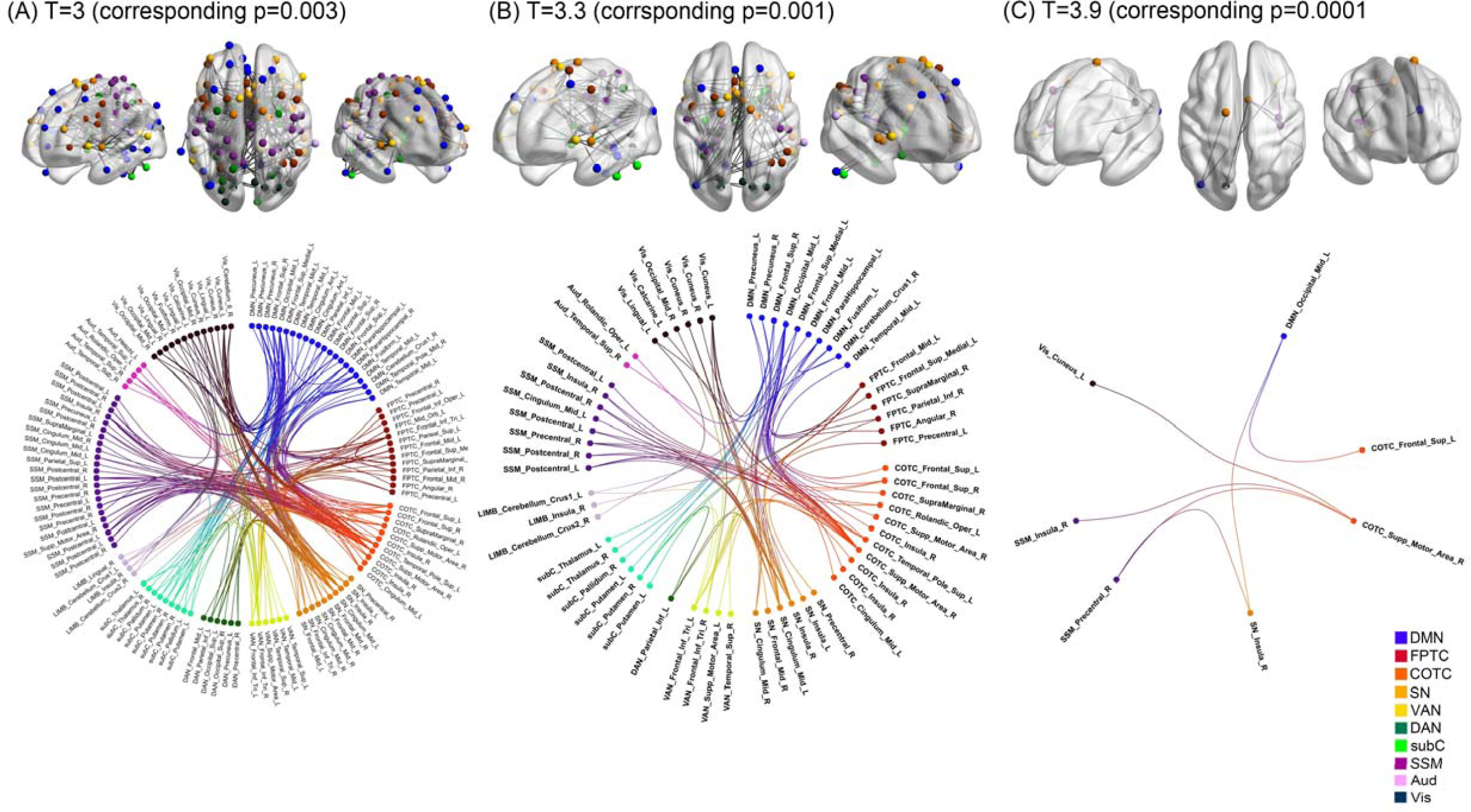
Group differences in functional connectivity based on different height thresholds in the network-based statistic (NBS). Across different t-statistics height thresholds (namely t=3 corresponding to uncorrected p=0.003; t=3.3 corresponding to uncorrected p=0.001; t=3.9 corresponding to uncorrected p=0.0001), the NBS consistently identified the similar patterns of hyperconnectivity between the DMN and attention and cognitive control networks, the DMN and subcortical regions, and the salience/cingulo-opercular network and sensory processing networks. These results were in line with the main discovered connectivity differences based on t=3.5 corresponding to uncorrected p=0.0005.

**Supplementary Figure 4.**
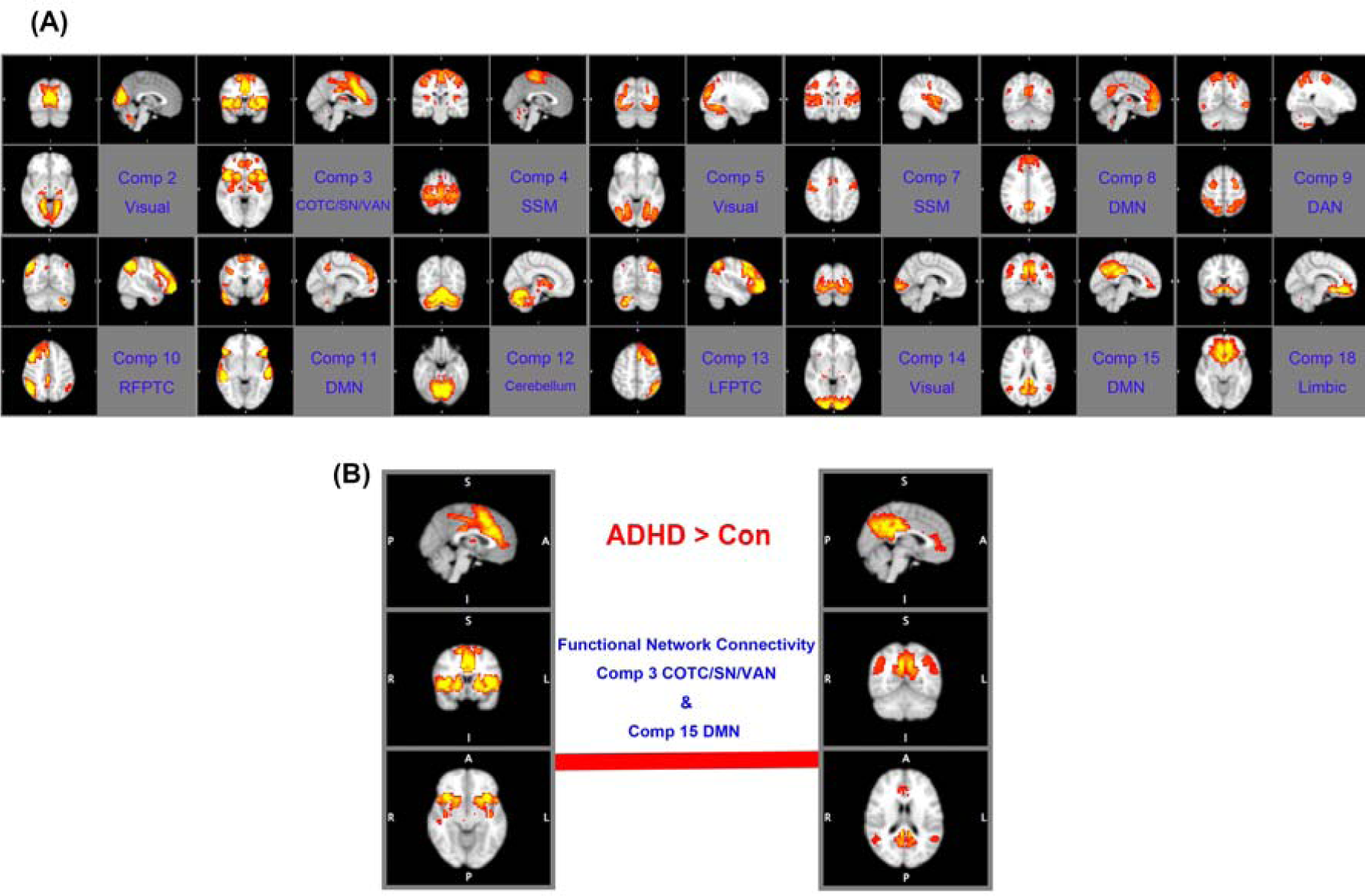
Independent component analysis (ICA) on neuroimaging data. (A) Based on the group ICA, we identified 20 spatial components. The topology of 14 components related to recognized functional brain networks (Yeo *et al.*, 2011; Cocchi *et al.*, 2012). These 14 components were used for the confirmatory functional network connectivity analysis (see text) (Jafri *et al.*, 2008). (B) Results from the functional network connectivity analysis are presented. Relative to the controls, adults with ADHD exhibited a significantly increased positive interaction between the default-mode and cingulo-opercular/salience networks (false discovery rate corrected *q*=0.044).

**Supplementary Figure 5.**
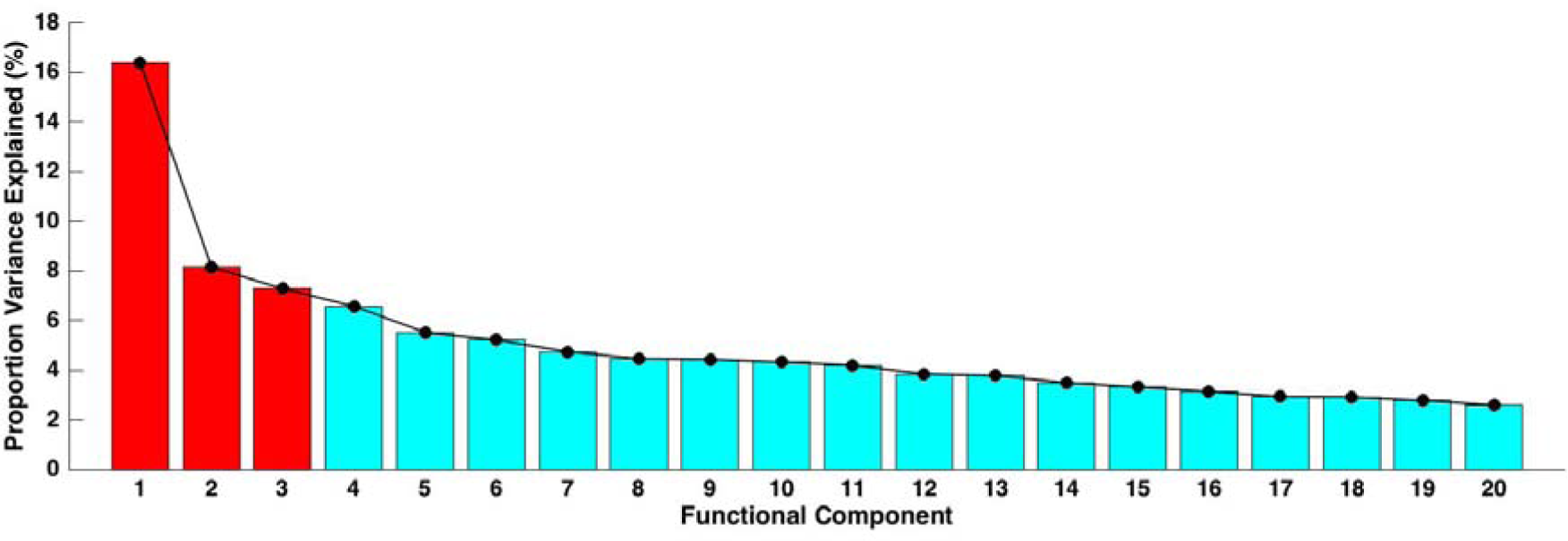
The proportion of variance explained by the eigenvectors defined by a principal component analysis on functional connectivity differences between ADHD and controls (derived from the network-based statistics). The three eigenvectors (red) used in the canonical correlation analysis (CCA, see text) explained 31.83% of the total variance in between-groups connectivity. Including two extra eigenvectors allows to explain 43.90% of the variance. CCA based on both three and five eigenvectors yielded a similarly significant CCA mode.

**Supplementary Figure 6.**
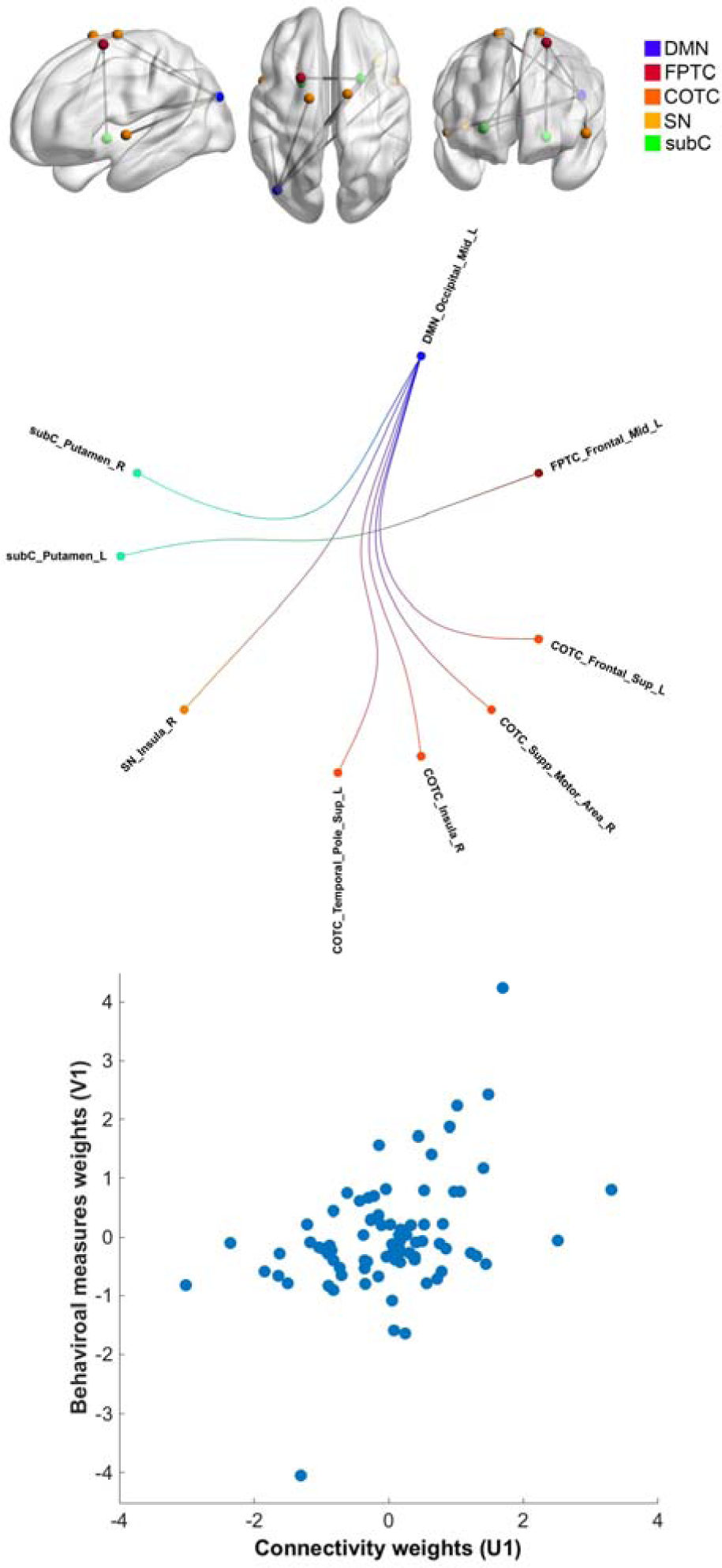
Supplementary canonical correlation analysis (CCA) based on the altered functional connectivity identified by the supplementary network-based statistic (NBS) adjusting for gender/sex, levels of in-scanner head motion, and age. This supplementary CCA yielded one significant mode, similar to the main result (Figure 3), which linked the brain connectivity and clinical symptoms-intelligence. The functional connections expressing the strongest positive associations in this mode from the supplementary analysis also implicated connectivity between the DMN and cingulo-opercular, as well as the DMN and subcortical regions.

**Supplementary Figure 7.**
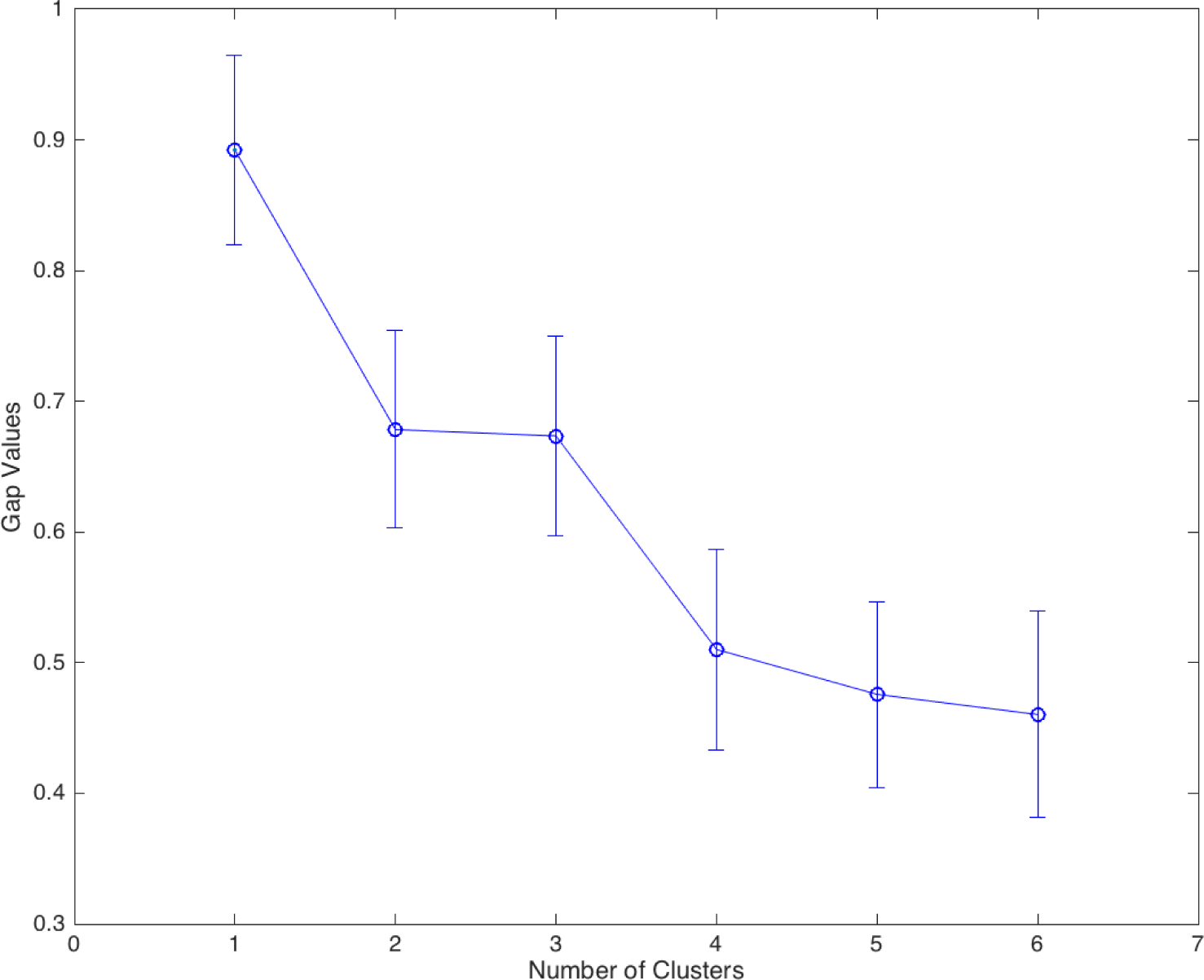
The gap statistic for the k-means clustering method (based on the feature vectors of individual participant’s weight derived from the connectivity and symptoms matrices of canonical correlation analysis). The gap statistic estimates the optimal number of clusters by searching the local maximum of the graph, then selecting the smallest *k* within one standard error (as indicated by the bars in the figure) of the local max [Gap(*k*) ≥ Gap(*k+1*) -*SE_k+1_*]. Based on the gap statistic, the suggested optimal number of clusters was one.

**Supplementary Figure 8.**
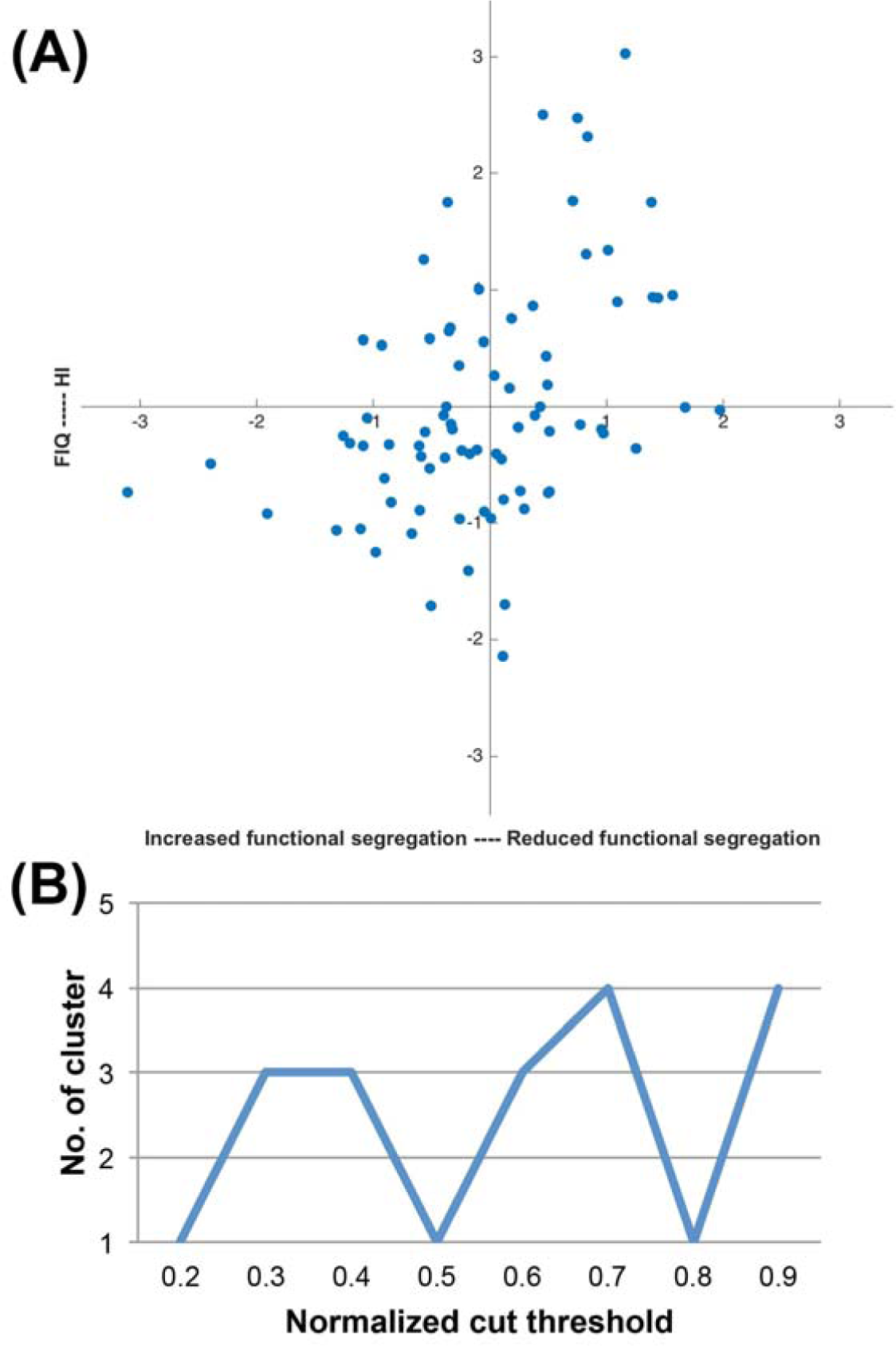
Test for ADHD categorical biotypes. (A) *K*-means analysis failed to reveal valid clusters based on the individual associations between functional connectivity and behavior. The absence of clear clusters in the data is evident from visual inspection of the figure. (B) The number (No.) of clusters detected by the multi-view spectral clustering algorithm changed as a function of the preset cut threshold, indicating that no stable decomposition was achievable. Overall, results from these analyses provide compelling evidence for the absence of non-overlapping clusters in the data. FIQ=full-scale IQ; HI=hyperactivity-impulsivity.

**Supplementary Figure 9.**
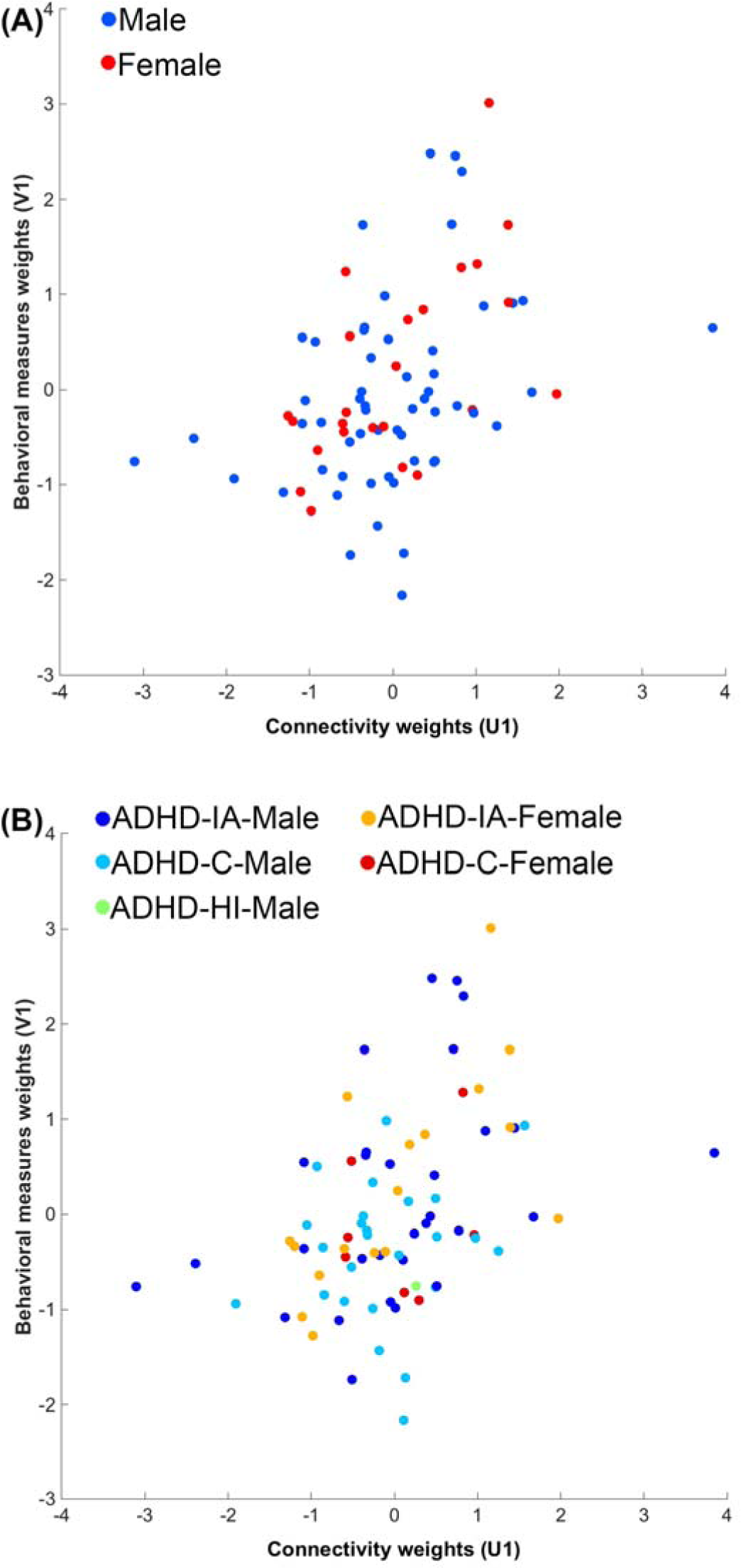
(A) Males and females with ADHD, (B) regardless of the clinical subtypes, were distributed evenly along the one-dimensional axis identified by the main CCA.

## References

Acosta, M. T., Castellanos, F. X., Bolton, K. L., Balog, J. Z., Eagen, P., Nee, L., Jones, J., Palacio, L., Sarampote, C., Russell, H. F., Berg, K., Arcos-Burgos, M. and Muenke, M. (2008). Latent class subtyping of attention-deficit/hyperactivity disorder and comorbid conditions. J Am Acad Child Adolesc Psychiatry 47, 797–807.

Asherson, P., Buitelaar, J., Faraone, S. V. and Rohde, L. A. (2016). Adult attention-deficit hyperactivity disorder: key conceptual issues. Lancet Psychiatry 3, 568–578.

Barber, A. D., Jacobson, L. A., Wexler, J. L., Nebel, M. B., Caffo, B. S., Pekar, J. J. and Mostofsky, S. H. (2015). Connectivity supporting attention in children with attention deficit hyperactivity disorder. Neuroimage Clin 7, 68–81.

Barch, D. M. (2017). Biotypes: Promise and Pitfalls. Biol Psychiatry 82, 2–3.

Birn, R. M., Molloy, E. K., Patriat, R., Parker, T., Meier, T. B., Kirk, G. R., Nair, V. A., Meyerand, M. E. and Prabhakaran, V. (2013). The effect of scan length on the reliability of resting-state fMRI connectivity estimates. Neuroimage 83, 550–558.

Cai, W., Chen, T., Szegletes, L., Supekar, K. and Menon, V. (2017). Aberrant Time-varying Cross-Network Interactions in Children with Attention-Deficit/Hyperactivity Disorder and Its Relation to Attention Deficits. Biological Psychiatry: Cognitive Neuroscience and Neuroimaging.

Castellanos, F. X. and Aoki, Y. (2016). Intrinsic Functional Connectivity in Attention-Deficit/Hyperactivity Disorder: A Science in Development. Biol Psychiatry Cogn Neurosci Neuroimaging 1, 253–261.

Cheung, C. H., Rijdijk, F., McLoughlin, G., Faraone, S. V., Asherson, P. and Kuntsi, J. (2015). Childhood predictors of adolescent and young adult outcome in ADHD. J Psychiatr Res 62, 92–100.

Clementz, B. A., Sweeney, J. A., Hamm, J. P., Ivleva, E. I., Ethridge, L. E., Pearlson, G. D., Keshavan, M. S. and Tamminga, C. A. (2016). Identification of Distinct Psychosis Biotypes Using Brain-Based Biomarkers. Am J Psychiatry 173, 373–384.

Cocchi, L., Bramati, I. E., Zalesky, A., Furukawa, E., Fontenelle, L. F., Moll, J., Tripp, G. and Mattos, P. (2012). Altered functional brain connectivity in a non-clinical sample of young adults with attention-deficit/hyperactivity disorder. J Neurosci 32, 17753–17761.

Cole, M. W., Ito, T., Bassett, D. S. and Schultz, D. H. (2016). Activity flow over resting-state networks shapes cognitive task activations. Nat Neurosci 19, 1718–1726.

Conners, C. K., D., E. and E., S. (1999). Conners’ adult ADHD rating scales (CAARS). MHS: New York.

Costa Dias, T. G., Iyer, S. P., Carpenter, S. D., Cary, R. P., Wilson, V. B., Mitchell, S. H., Nigg, J. T. and Fair, D. A. (2015). Characterizing heterogeneity in children with and without ADHD based on reward system connectivity. Dev Cogn Neurosci 11, 155–174.

Cuthbert, B. N. (2015). Research Domain Criteria: toward future psychiatric nosologies. Dialogues Clin Neurosci 17, 89–97.

Demontis, D., Walters, R. K., Martin, J., Mattheisen, M., Als, T. D., Agerbo, E., Belliveau, R., Bybjerg-Grauholm, J., Bækved-Hansen, M. and Cerrato, F. (2017). Discovery Of The First Genome-Wide Significant Risk Loci For ADHD. bioRxiv, 145581.

Drysdale, A. T., Grosenick, L., Downar, J., Dunlop, K., Mansouri, F., Meng, Y., Fetcho, R. N., Zebley, B., Oathes, D. J., Etkin, A., Schatzberg, A. F., Sudheimer, K., Keller, J., Mayberg, H. S., Gunning, F. M., Alexopoulos, G. S., Fox, M. D., Pascual-Leone, A., Voss, H. U., Casey, B. J., Dubin, M. J. and Liston, C. (2017). Resting-state connectivity biomarkers define neurophysiological subtypes of depression. Nat Med 23, 28–38.

Fair, D. A., Bathula, D., Nikolas, M. A. and Nigg, J. T. (2012). Distinct neuropsychological subgroups in typically developing youth inform heterogeneity in children with ADHD. Proc Natl Acad Sci U S A 109, 6769–6774.

Faraone, S. V. and Biederman, J. (2016). Can Attention-Deficit/Hyperactivity Disorder Onset Occur in Adulthood? JAMA Psychiatry 73, 655–656.

Gallo, E. F. and Posner, J. (2016). Moving towards causality in attention-deficit hyperactivity disorder: overview of neural and genetic mechanisms. Lancet Psychiatry 3, 555–567.

Gates, K. M., Molenaar, P. C., Iyer, S. P., Nigg, J. T. and Fair, D. A. (2014). Organizing heterogeneous samples using community detection of GIMME-derived resting state functional networks. PLoS One 9, e91322.

Hearne, L. J., Mattingley, J. B. and Cocchi, L. (2016). Functional brain networks related to individual differences in human intelligence at rest. Sci Rep 6, 32328.

Hennig, C., Meila, M., Murtagh, F. and Rocci, R. (2015). Handbook ofCluster Analysis. CRC Press.

Hoogman, M., Bralten, J., Hibar, D. P., Mennes, M., Zwiers, M. P., Schweren, L. S., van Hulzen, K. J., Medland, S. E., Shumskaya, E., Jahanshad, N., Zeeuw, P., Szekely, E., Sudre, G., Wolfers, T., Onnink, A. M., Dammers, J. T., Mostert, J. C., Vives-Gilabert, Y., Kohls, G., Oberwelland, E., Seitz, J., Schulte-Ruther, M., Ambrosino, S., Doyle, A. E., Hovik, M. F., Dramsdahl, M., Tamm, L., van Erp, T. G., Dale, A., Schork, A., Conzelmann, A., Zierhut, K., Baur, R., McCarthy, H., Yoncheva, Y. N., Cubillo, A., Chantiluke, K., Mehta, M. A., Paloyelis, Y., Hohmann, S., Baumeister, S., Bramati, I., Mattos, P., Tovar-Moll, F., Douglas, P., Banaschewski, T., Brandeis, D., Kuntsi, J., Asherson, P., Rubia, K., Kelly, C., Martino, A. D., Milham, M. P., Castellanos, F. X., Frodl, T., Zentis, M., Lesch, K. P., Reif, A., Pauli, P., Jernigan, T. L., Haavik, J., Plessen, K. J., Lundervold, A. J., Hugdahl, K., Seidman, L. J., Biederman, J., Rommelse, N., Heslenfeld, D. J., Hartman, C. A., Hoekstra, P. J., Oosterlaan, J., Polier, G. V., Konrad, K., Vilarroya, O., Ramos-Quiroga, J. A., Soliva, J. C., Durston, S., Buitelaar, J. K., Faraone, S. V., Shaw, P., Thompson, P. M. and Franke, B. (2017). Subcortical brain volume differences in participants with attention deficit hyperactivity disorder in children and adults: a cross-sectional mega-analysis. Lancet Psychiatry 4, 310–319.

Jafri, M. J., Pearlson, G. D., Stevens, M. and Calhoun, V. D. (2008). A method for functional network connectivity among spatially independent resting-state components in schizophrenia. Neuroimage 39, 1666–1681.

Kaczkurkin, A. N., Moore, T. M., Calkins, M. E., Ciric, R., Detre, J. A., Elliott, M. A., Foa, E. B., Garcia de la Garza, A., Roalf, D. R., Rosen, A., Ruparel, K., Shinohara, R. T., Xia, C. H., Wolf, D. H., Gur, R. E., Gur, R. C. and Satterthwaite, T. D. (2017). Common and dissociable regional cerebral blood flow differences associate with dimensions of psychopathology across categorical diagnoses. Mol Psychiatry.

Karalunas, S. L., Fair, D., Musser, E. D., Aykes, K., Iyer, S. P. and Nigg, J. T. (2014). Subtyping attention-deficit/hyperactivity disorder using temperament dimensions: toward biologically based nosologic criteria. JAMA Psychiatry 71, 1015–1024.

Keyes, K. M., Platt, J., Kaufman, A. S. and McLaughlin, K. A. (2017). Association of Fluid Intelligence and Psychiatric Disorders in a Population-Representative Sample of US Adolescents. JAMA Psychiatry 74, 179–188.

Krzanowski, W. (2000). Principles of multivariate analysis. OUP Oxford.

Kumar, A. and Daumé, H. (2011). A co-training approach for multi-view spectral clustering. In Proceedings of the 28th International Conference on Machine Learning (ICML-11)pp. 393–400.

Kundu, P., Inati, S. J., Evans, J. W., Luh, W. M. and Bandettini, P. A. (2012). Differentiating BOLD and non-BOLD signals in fMRI time series using multi-echo EPI. Neuroimage 60, 1759–1770.

Kundu, P., Voon, V., Balchandani, P., Lombardo, M. V., Poser, B. A. and Bandettini, P. (2017). Multi-Echo fMRI: A Review of Applications in fMRI Denoising and Analysis of BOLD Signals. Neuroimage.

Lin, H. Y. and Gau, S. S. (2015). Atomoxetine Treatment Strengthens an Anti-Correlated Relationship between Functional Brain Networks in Medication-Naive Adults with Attention-Deficit Hyperactivity Disorder: A Randomized Double-Blind Placebo-Controlled Clinical Trial. Int J Neuropsychopharmacol 19, pyv094.

Lin, Y. J., Yang, L. K. and Gau, S. S. (2016). Psychiatric comorbidities of adults with early- and late-onset attention-deficit/hyperactivity disorder. Aust N Z J Psychiatry 50, 548–556.

Lombardo, M. V., Auyeung, B., Holt, R. J., Waldman, J., Ruigrok, A. N., Mooney, N., Bullmore, E. T., Baron-Cohen, S. and Kundu, P. (2016). Improving effect size estimation and statistical power with multi-echo fMRI and its impact on understanding the neural systems supporting mentalizing. Neuroimage 142, 55–66.

Marcus, D. K. and Barry, T. D. (2011). Does attention-deficit/hyperactivity disorder have a dimensional latent structure? A taxometric analysis. J Abnorm Psychol 120, 427–442.

Marquand, A. F., Wolfers, T., Mennes, M., Buitelaar, J. and Beckmann, C. F. (2016). Beyond Lumping and Splitting: A Review of Computational Approaches for Stratifying Psychiatric Disorders. Biol Psychiatry Cogn Neurosci Neuroimaging 1, 433–447.

Menon, V. (2011). Large-scale brain networks and psychopathology: a unifying triple network model. Trends Cogn Sci 15, 483–506.

Mostert, J. C., Hoogman, M., Onnink, A. M., van Rooij, D., von Rhein, D., van Hulzen, K. J., Dammers, J., Kan, C. C., Buitelaar, J. K., Norris, D. G. and Franke, B. (2015). Similar Subgroups Based on Cognitive Performance Parse Heterogeneity in Adults With ADHD and Healthy Controls. J Atten Disord.

Perry, A., Wen, W., Kochan, N. A., Thalamuthu, A., Sachdev, P. S. and Breakspear, M. (2017). The independent influences of age and education on functional brain networks and cognition in healthy older adults. Hum Brain Mapp 38, 5094–5114.

Power, J. D., Cohen, A. L., Nelson, S. M., Wig, G. S., Barnes, K. A., Church, J. A., Vogel, A. C., Laumann, T. O., Miezin, F. M., Schlaggar, B. L. and Petersen, S. E. (2011). Functional network organization of the human brain. Neuron 72, 665–678.

Rommelse, N., van der Kruijs, M., Damhuis, J., Hoek, I., Smeets, S., Antshel, K. M., Hoogeveen, L. and Faraone, S. V. (2016). An evidenced-based perspective on the validity of attention-deficit/hyperactivity disorder in the context of high intelligence. Neurosci Biobehav Rev 71, 21–47.

Schnack, H. G. and Kahn, R. S. (2016). Detecting Neuroimaging Biomarkers for Psychiatric Disorders: Sample Size Matters. Front Psychiatry 7, 50.

Smith, S. M., Nichols, T. E., Vidaurre, D., Winkler, A. M., Behrens, T. E., Glasser, M. F., Ugurbil, K., Barch, D. M., van Essen, D. C. and Miller, K. L. (2015). A positive-negative mode of population covariation links brain connectivity, demographics and behavior. Nat Neurosci 18, 1565–1567.

Tibshirani, R., Walther, G. and Hastie, T. (2001). Estimating the number of clusters in a data set via the gap statistic. Journal of the Royal Statistical Society: Series B (Statistical Methodology) 63, 411–423.

van Dijk, K. R., Hedden, T., Venkataraman, A., Evans, K. C., Lazar, S. W. and Buckner, R. L. (2010). Intrinsic functional connectivity as a tool for human connectomics: theory, properties, and optimization. J Neurophysiol 103, 297–321.

van Rooij, D., Hartman, C. A., Mennes, M., Oosterlaan, J., Franke, B., Rommelse, N., Heslenfeld, D., Faraone, S. V., Buitelaar, J. K. and Hoekstra, P. J. (2015). Altered neural connectivity during response inhibition in adolescents with attention-deficit/hyperactivity disorder and their unaffected siblings. Neuroimage Clin 7, 325–335.

Wechsler, D. (1997). Wechsler Adult Intelligence Scale - Third Edition (WAISIII). Psychological Corporation: San Antonio, TX

Willcutt, E. G., Nigg, J. T., Pennington, B. F., Solanto, M. V., Rohde, L. A., Tannock, R., Loo, S. K., Carlson, C. L., McBurnett, K. and Lahey, B. B. (2012). Validity of DSM-IV attention deficit/hyperactivity disorder symptom dimensions and subtypes. J Abnorm Psychol 121, 991–1010.

Yeh, C. B., Gau, S. S., Kessler, R. C. and Wu, Y. Y. (2008). Psychometric properties of the Chinese version of the adult ADHD Self-report Scale. Int J Methods Psychiatr Res 17, 45–54.

Zalesky, A., Fornito, A. and Bullmore, E. T. (2010). Network-based statistic: identifying differences in brain networks. Neuroimage 53, 1197–1207.

## References

Abdi, H. and Williams, L. J. (2010). Principal component analysis. Wiley interdisciplinary reviews: computational statistics 2, 433–459.

Beckmann, C. F. and Smith, S. M. (2004). Probabilistic independent component analysis for functional magnetic resonance imaging. IEEE Trans Med Imaging 23, 137–152.

Benjamini, Y. and Hochberg, Y. (1995). Controlling the false discovery rate: a practical and powerful approach to multiple testing. Journal of the royal statistical society. Series B (Methodological), 289–300.

Chang, L. R., Chiu, Y. N., Wu, Y. Y. and Gau, S. S. (2013). Father’s parenting and father-child relationship among children and adolescents with attention-deficit/hyperactivity disorder. Compr Psychiatry 54, 128–140.

Chao, C. Y., Gau, S. S., Mao, W. C., Shyu, J. F., Chen, Y. C. and Yeh, C. B. (2008). Relationship of attention-deficit-hyperactivity disorder symptoms, depressive/anxiety symptoms, and life quality in young men. Psychiatry Clin Neurosci 62, 421–426.

Chen, H., Li, K., Zhu, D., Jiang, X., Yuan, Y., Lv, P., Zhang, T., Guo, L., Shen, D. and Liu, T. (2013). Inferring group-wise consistent multimodal brain networks via multi-view spectral clustering. IEEE Trans Med Imaging 32, 1576–1586.

Cocchi, L., Harrison, B. J., Pujol, J., Harding, I. H., Fornito, A., Pantelis, C. and Yucel, M. (2012). Functional alterations of large-scale brain networks related to cognitive control in obsessive-compulsive disorder. Hum Brain Mapp 33, 1089–1106.

Craddock, R. C., James, G. A., Holtzheimer, P. E., 3rd, Hu, X. P. and Mayberg, H. S. (2012). A whole brain fMRI atlas generated via spatially constrained spectral clustering. Hum Brain Mapp 33, 1914–1928.

DiStefano, C., Zhu, M. and Mindrila, D. (2009). Understanding and using factor scores: Considerations for the applied researcher. Practical Assessment, Research & Evaluation 14, 1–11.

Dolnicar, S. (2002). A review of unquestioned standards in using cluster analysis for data-driven market segmentation. In CD Conference Proceedings of the Australian and New Zealand Marketing Academy Conference 2002 (ANZMAC 2002): Deakin University, Melbourne.

Dosenbach, N. U., Nardos, B., Cohen, A. L., Fair, D. A., Power, J. D., Church, J. A., Nelson, S. M., Wig, G. S., Vogel, A. C., Lessov-Schlaggar, C. N., Barnes, K. A., Dubis, J. W., Feczko, E., Coalson, R. S., Pruett, J. R., Jr., Barch, D. M., Petersen, S. E. and Schlaggar, B. L. (2010). Prediction of individual brain maturity using fMRI. Science 329, 1358–1361.

Gau, S. F. and Soong, W. T. (1999). Psychiatric comorbidity of adolescents with sleep terrors or sleepwalking: a case-control study. Aust N Z J Psychiatry 33, 734–739.

Gau, S. S., Chong, M. Y., Chen, T. H. and Cheng, A. T. (2005). A 3-year panel study of mental disorders among adolescents in Taiwan. Am J Psychiatry 162, 1344–1350.

Gau, S. S., Kessler, R. C., Tseng, W. L., Wu, Y. Y., Chiu, Y. N., Yeh, C. B. and Hwu, H. G. (2007). Association between sleep problems and symptoms of attention-deficit/hyperactivity disorder in young adults. Sleep 30, 195–201.

Gau, S. S., Shang, C. Y., Liu, S. K., Lin, C. H., Swanson, J. M., Liu, Y. C. and Tu, C. L. (2008). Psychometric properties of the Chinese version of the Swanson, Nolan, and Pelham, version IV scale - parent form. Int J Methods Psychiatr Res 17, 35–44.

Hennig, C. (2008). Dissolution point and isolation robustness: robustness criteria for general cluster analysis methods. Journal of multivariate analysis 99, 1154–1176.

Huettel, S. A., Song, A. W. and McCarthy, G. (2008). Functional magnetic resonance imaging. Sinauer Associates: Sunderland.

Hyvarinen, A. (1999). Fast and robust fixed-point algorithms for independent component analysis. IEEE Trans Neural Netw 10, 626–634.

Kononenko, I. and Kukar, M. (2007). Machine learning and data mining: introduction to principles and algorithms. Horwood Publishing.

Kumar, A. and Daumé, H. (2011). A co-training approach for multi-view spectral clustering. In Proceedings of the 28th International Conference on Machine Learning (ICML-11) pp. 393–400.

Kundu, P., Brenowitz, N. D., Voon, V., Worbe, Y., Vertes, P. E., Inati, S. J., Saad, Z. S., Bandettini, P. A. and Bullmore, E. T. (2013). Integrated strategy for improving functional connectivity mapping using multiecho fMRI. Proc Natl Acad Sci U S A 110, 16187–16192.

Lv, J., Iraji, A., Ge, F., Zhao, S., Hu, X., Zhang, T., Han, J., Guo, L., Kou, Z. and Liu, T. (2016). Temporal Concatenated Sparse Coding of Resting State fMRI Data Reveal Network Interaction Changes in mTBI. In International Conference on Medical Image Computing and Computer-Assisted Interventionpp. 46–54. Springer.

Ni, H. C., Lin, Y. J., Gau, S. S., Huang, H. C. and Yang, L. K. (2013a). An Open-Label, Randomized Trial of Methylphenidate and Atomoxetine Treatment in Adults With ADHD. J Atten Disord.

Ni, H. C., Shang, C. Y., Gau, S. S., Lin, Y. J., Huang, H. C. and Yang, L. K. (2013b). A head-to-head randomized clinical trial of methylphenidate and atomoxetine treatment for executive function in adults with attention-deficit hyperactivity disorder. Int J Neuropsychopharmacol 16, 1959–1973.

Orvaschel, H., Puig-Antich, J., Chambers, W., Tabrizi, M. A. and Johnson, R. (1982). Retrospective assessment of prepubertal major depression with the Kiddie-SADS-e. J Am Acad Child Psychiatry 21, 392–397.

Shi, J. and Malik, J. (2000). Normalized cuts and image segmentation. IEEE Transactions on pattern analysis and machine intelligence 22, 888–905.

Smith, S. M., Fox, P. T., Miller, K. L., Glahn, D. C., Fox, P. M., Mackay, C. E., Filippini, N., Watkins, K. E., Toro, R., Laird, A. R. and Beckmann, C. F. (2009). Correspondence of the brain’s functional architecture during activation and rest. Proc Natl Acad Sci U S A 106, 13040–13045.

Smith, S. M., Hyvarinen, A., Varoquaux, G., Miller, K. L. and Beckmann, C. F. (2014). Group-PCA for very large fMRI datasets. Neuroimage 101, 738–749.

Smith, S. M., Nichols, T. E., Vidaurre, D., Winkler, A. M., Behrens, T. E., Glasser, M. F., Ugurbil, K., Barch, D. M., Van Essen, D. C. and Miller, K. L. (2015). A positive-negative mode of population covariation links brain connectivity, demographics and behavior. Nat Neurosci 18, 1565–1567.

Swanson, J. M., Kraemer, H. C., Hinshaw, S. P., Arnold, L. E., Conners, C. K., Abikoff, H. B., Clevenger, W., Davies, M., Elliott, G. R., Greenhill, L. L., Hechtman, L., Hoza, B., Jensen, P. S., March, J. S., Newcorn, J. H., Owens, E. B., Pelham, W. E., Schiller, E., Severe, J. B., Simpson, S., Vitiello, B., Wells, K., Wigal, T. and Wu, M. (2001). Clinical relevance of the primary findings of the MTA: success rates based on severity of ADHD and ODD symptoms at the end of treatment. J Am Acad Child Adolesc Psychiatry 40, 168–179.

Takahashi, M., Goto, T., Takita, Y., Chung, S. K., Wang, Y. and Gau, S. S. (2014). Open-label, dose-titration tolerability study of atomoxetine hydrochloride in Korean, Chinese, and Taiwanese adults with attention-deficit/hyperactivity disorder. Asia Pac Psychiatry 6, 62–70.

Tao, H., Hou, C. and Yi, D. (2014). Multiple-view spectral embedded clustering using a co-training approach. In Computer Engineering and Networkingpp. 979–987. Springer.

Tibshirani, R., Walther, G. and Hastie, T. (2001). Estimating the number of clusters in a data set via the gap statistic. Journal ofthe Royal Statistical Society: Series B (Statistical Methodology) 63, 411–423.

Venkataraman, A., Van Dijk, K. R., Buckner, R. L. and Golland, P. (2009). Exploring Functional Connectivity in Fmri Via Clustering. Proc IEEE Int Conf Acoust Speech Signal Process 2009, 441–444.

Yang, H. N., Tai, Y. M., Yang, L. K. and Gau, S. S. (2013). Prediction of childhood ADHD symptoms to quality of life in young adults: adult ADHD and anxiety/depression as mediators. Res Dev Disabil 34, 3168–3181.

Yeo, B. T., Krienen, F. M., Sepulcre, J., Sabuncu, M. R., Lashkari, D., Hollinshead, M., Roffman, J. L., Smoller, J. W., Zollei, L., Polimeni, J. R., Fischl, B., Liu, H. and Buckner, R. L. (2011). The organization of the human cerebral cortex estimated by intrinsic functional connectivity. J Neurophysiol 106, 1125–1165.

